# Identification Drug Targets for Oxaliplatin-Induced Cardiotoxicity without Affecting Cancer Treatment through Inter Variability Cross-Correlation Analysis (IVCCA)

**DOI:** 10.1101/2024.02.11.579390

**Authors:** Junwei Du, Leland C. Sudlow, Hridoy Biswas, Joshua D. Mitchell, Shamim Mollah, Mikhail Y. Berezin

## Abstract

The successful treatment of side effects of chemotherapy faces two major limitations: the need to avoid interfering with pathways essential for the cancer-destroying effects of the chemotherapy drug, and the need to avoid helping tumor progression through cancer promoting cellular pathways. To address these questions and identify new pathways and targets that satisfy these limitations, we have developed the bioinformatics tool Inter Variability Cross-Correlation Analysis (IVCCA). This tool calculates the cross-correlation of differentially expressed genes, analyzes their clusters, and compares them across a vast number of known pathways to identify the most relevant target(s). To demonstrate the utility of IVCCA, we applied this platform to RNA-seq data obtained from the hearts of the animal models with oxaliplatin-induced CTX. RNA-seq of the heart tissue from oxaliplatin treated mice identified 1744 differentially expressed genes with False Discovery Rate (FDR) less than 0.05 and fold change above 1.5 across nine samples. We compared the results against traditional gene enrichment analysis methods, revealing that IVCCA identified additional pathways potentially involved in CTX beyond those detected by conventional approaches. The newly identified pathways such as energy metabolism and several others represent promising target for therapeutic intervention against CTX, while preserving the efficacy of the chemotherapy treatment and avoiding tumor proliferation. Targeting these pathways is expected to mitigate the damaging effects of chemotherapy on cardiac tissues and improve patient outcomes by reducing the incidence of heart failure and other cardiovascular complications, ultimately enabling patients to complete their full course of chemotherapy with improved quality of life and survival rates.

## INTRODUCTION

Cardiac dysfunction resulting from exposure to cancer therapeutics was first recognized in the 1960s (1). Heart failure associated with certain cancer drugs, such as the anthracyclines, was soon recognized as an important side effect. It has also been observed that not all cancer treatments affect the heart in the same way and therefore these agents cannot be viewed as a single class of drugs (2). Each class of drugs requires an individual approach. Since then, significant efforts have been undertaken to recognize the signs of chemotherapy induced cardiotoxicity (CTX) early on and to identify methods to mitigate CTX without compromising the cancer treatment. Clinical data on cardiotoxicities reported during chemotherapy reveal that cardiac damages are associated with many pathways that include oxidative stress, mitochondrial dysfunction, apoptosis, inflammation, and damage to the myocardium (3,4).

Chemotherapies for cancer treatment cause devastating adverse side effects, leading to significantly increased morbidity and mortality among cancer patients. With a larger number of cancer survivors, these side effects have become a critical and escalating concern in oncology, leading to significantly increased morbidity and mortality among cancer patients (5,6). Many chemotherapies agents induce a range of cardiac complications such as arrhythmias, myocardial ischemia, thromboembolic disease, left ventricular dysfunction and heart failure (7–9). The risk of cardiotoxicity is influenced by several factors including the type and dose of chemotherapy, patient’s age, pre-existing cardiovascular disease, other comorbidities, and concurrent cardiovascular risk factors. Importantly, this increased risk persists even after the termination of chemotherapy, with late-onset CTX being a grave concern. The development of CTX can significantly impact the quality of life of survivors and may, in severe cases, become life-threatening. There is an urgent need for cardiac monitoring during chemotherapy, along with the development of strategies to predict prevent and treat this serious side effect (10).

While some mechanisms for CTX with some drugs (i.e., anthracyclines) have been proposed (11) the mechanisms of CTX for many drugs are not yet developed. Oxaliplatin is the first line of defense in colorectal cancers and is used against other malignancies, including gastric, pancreatic, and advanced hepatocellular carcinomas. Oxaliplatin’s efficacy is limited by its off-target toxicity such as peripheral neuropathy, nephrotoxicity, gastrointestinal toxicity, and others (12–17). One of the typical, but less studied side effects of oxaliplatin is its adverse effect on the heart. Patients treated with oxaliplatin often experience rapid breathing, chest pain, tachycardia, and arrhythmias. Although emergency situations in oxaliplatin-treated patients are relatively rare compared to other drugs like anthracyclines (18,19) or 5-fluorouacil (20), it is a rising concern given the increasing number of patients treated with oxaliplatin alone or in combination with other drugs. A growing number of cases related to severe coronary and cardiotoxicity of oxaliplatin alone (21,22) or together with 5-fluoruracil or FOLFOX (23–26) have been reported. Understanding the mechanism of oxaliplatin-induced CTX to improve prediction and treat is of the paramount importance.

Identification of promising drug targets for drug discovery relies on the utilization of omics tools, with RNA-seq being one of the most commonly used techniques. This method enables the identification of differentially expressed genes (DEGs) between experimental groups, such as diseased and healthy individuals or treated and untreated animal models. To make sense of the large amount of data generated by RNA-seq experiments, bioinformatics tools are applied to analyze gene expression patterns, identify enriched gene ontology terms, and perform pathway analysis. They provide statistical methods and algorithms to assess the significance of differential expression, highlighting genes and pathways that play crucial roles in disease development or treatment response. However, there is a lack of efficient tools to discern promising drug targets and pathways without being confounded by other pathways that may negatively interact with treatment strategies, such as chemotherapy. Establishing a treatment for the side effects of chemotherapy without compromising the efficacy of the cancer treatment itself is akin to navigating between mythical Scylla and Charybdis. It requires a precise approach, where one must carefully avoid several equally dangerous hazards to reach the desired therapeutic effect. There is a clear need for new and improved tools that can better navigate the complexity of gene expression data and pathway interactions in the context of drug discovery and development.

Addressing this challenge in today’s oncology, we have developed the Inter Variability Cross-Correlation Analysis (IVCCA) method. This method facilitates the ranking of pathways and aids in identifying significant pathways and genes, based on the individual expression of each differentially expressed gene, as illustrated in **Figure 1**. This approach enables the determination of whether a pathway or gene with potential therapeutic value can be targeted safely, without adversely affecting chemotherapy or contributing to cancer growth. To explore this concept, we applied IVCCA to identify promising pathways and targets for treatment of oxaliplatin induced CTX.

**Figure 1.**
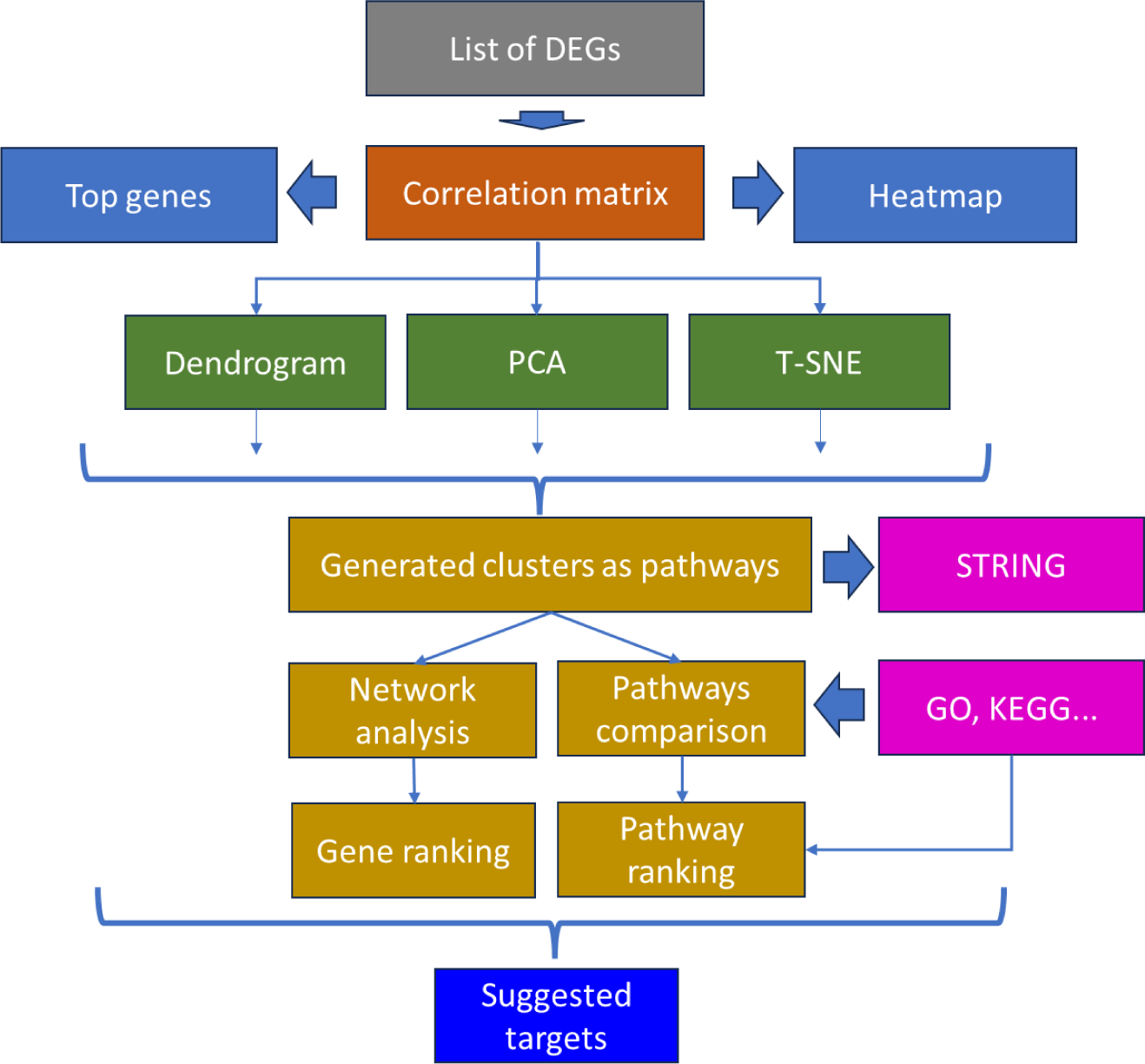
A general outline of the proposed pipeline analysis. Data is processed through a traditional pipeline of RNA-seq data preprocessing and differential expression genes (DEGs) extraction using specific filter parameters such as False Discovery Rate (FDR) <0.05 and fold change (FC) >1.5. The data are utilized to construct a correlation matrix, its correlation heatmap is generated to visualize DEGs’ correlation distribution. For further analysis, the absolute values of the correlations are ordered. The sorted heatmap aids in the visualization of the top genes. Clustering is performed using Dendrogram, Principal Component Analysis (PCA), and t-distributed Stochastic Neighbor Embedding (t-SNE) methods, followed by distance thresholding (for the Dendrogram results) or K-means (for the PCA and t-SNE results) for finer clustering. Clusters were analyzed via STRING (27) or network analysis to identify potential target genes. All pathways, including generated and existing ones from databases like GO and KEGG, are quantitatively compared using novel indices and ranked for relevance.

## METHODS

### RNA sequencing datasets

RNA-seq data were used from our previously published study (GEO repository GSE233805). Genes were considered to be differentially expressed with FDR<0.05 (28), and absolute fold change FC >1.5 or FC<-1.5. From 13775 genes identified in the dataset, 1744 DEGs based on these criteria were selected. The list of these genes and their expressions for individual mice are available in the Supplementary Information section in the Excel format (*RNA_seq_DEGs.xlsx*). A list of 1744 randomly selected genes was constructed from the same parental dataset of 13775 genes with neither FDR nor FC thresholds applied.

The list of genes from the selected pathways were downloaded from GO and KEGG. Some of the lists were generated by us based on literature data (Pubmed), publicly available websites (i.e., ECM pathways from the Matrisome Project (29) or based on our previous analysis (i.e., energy metabolic pathway (30)). The lists of these pathway specific genes are available in the Supplementary Information section as .*txt* files.

### Overall design of the Inter-Variability Cross-Correlation Analysis (IVCCA) GUI

We used the IVCCA graphic user interface in MATLAB to perform correlation analysis. Examples of the results of the analysis are presented in **Figure S1**. The correlation analyses were performed on the gene expression data (.xlsx, .cvs, or .tsv file formats). We ranked the genes based on the magnitudes of the absolute values of the correlation coefficients from all genes in the matrix normalized to the number of genes. We then performed clustering analyses using Dendrogram (by selecting a distance threshold), Principal Component Analysis (PCA) or t-distributed Stochastic Neighbor Embedding (t-SNE) analyses (via K-means testing). The optimal number of genes used in the clustering analyses was determined via elbow (31) and a silhouette (32) methods. Statistical metrics produced by IVCCA identified the most relevant pathways from large pathway databases, such as KEGG and GO, and aided in the identification of hub genes. The results of the different clustering and pathway analyses were compared cosine similarity matrix. We then used results of the correlation analyses and statistical metrics to perform network analyses to determine the connections between genes and identify genes with the highest or lowest connections as potential targets.

### Theory

The theory behind each of the algorithms is provided in the Supplemental Information as well as in the MATLAB package that can be downloaded from the GitHub developer platform: https://github.com/MikhailBerezin/IVCCA/.

### Dendrogram and Hierarchical clustering

For dendrogram and hierarchical clustering we used an Unweighted Pair Group Method with Arithmetic Mean (UPGMA) algorithm that builds a dendrogram by successively merging clusters based on the mean distances between their members (33). The code calculates the pairwise Euclidean distances between data points in the correlation matrix using the ‘*pdist*’ function implemented in MATLAB. Hierarchical clustering was then performed on the distance matrix using the ‘*average*’ linkage method. Genes with greater similarity in the Pearson correlation analysis are positioned closer together on the dendrogram. The code prompts the user to enter a color threshold. A user determined color threshold was used in to plotting the different clusters and a gene list was generated for each cluster.

### Gene-to-Pathways analysis

The correlation between a specified single gene and a set of genes in a given pathway or a cluster was examined. This step in the analysis involved calculating the average absolute value of the correlation between the gene of interest and that of each gene in the pathway. The correlations were visually presented, where positive correlations are shown in blue and negative correlations in red as shown in the example in **Figure S2**. We also calculated the average of the absolute values of these correlation coefficients, providing a summary statistic of the overall correlation of the single gene with the genes in a certain pathway.

### Compare Pathways analysis

We applied a cosine similarity function in IVCCA to calculate the difference between two groups of genes. The algorithm found genes that overlapped between a reference pathway and one or multiple sets of genes by calculating the cosine similarity for each providing a theoretical score between 0 and 1. Several indices were calculated. The Pathway Activated Index, (*PAI* Eq. 9, Supplemental Information) reflects the percentage of the genes found in the set to the total number of genes in the pathway. The Pathway Correlation Indices (*PCI_A* and *PCI_B*) which represents the average cross-correlations of gene expressions within a specific pathway or set of genes (*PCI_A*, Eq. 5, Supplemental Information) and across the entire dataset (*PCI_B*, Eq.5 Supplemental Information). The difference between the *PCI_A* and *PCI_B* is shown in **Figure S2**. Greater similarity between pathways provides values closer to 1. In practice, within our dataset for KEGG and GO pathways we observed values for *PCI_A* between 0.31 to 1 and for *PCI_B* from 0.44 to 0.8. The mathematical description of this process is given in Supplemental Information. Correlation-Expression Composite Index (*CECI*) represents the strength of the pathway and numerically equals to the product of *PAI* and *PCI_B*. The values of *CECI* for selected pathway are graphically presented as bar graphs in the descending order.

*Z-score* (Eq. 11, Supplemental Information) indicates how many standard deviations an element is from the mean and is directly proportional to *CECI*. *Z-score* determines statistical significance of each pathway. Pathways with the *Z_score* > *Z_score_critical* (where *Z_score_critical* = 1.96, see Eq. 17, Supplementary Information) were considered statistically.

The each analysis report contained the correlation data and gene names from the correlation table, presents the total number of genes in each pathway, the number of genes found in the current set, and several indices: Pathway Activated Index, (*PAI*), Pathway Correlation Index within the pathway *(PCI_A*), Pathway Correlation Index within the entire dataset, (*PCI_B*), Correlation-Expression Composite Index (*CECI*) and Z-score for that combination of gene(s) and pathway.

### Pathway enrichment analysis using established methods

The pathway enrichment analysis evaluated whether the DEGs were overrepresented in the pathway compared to what would be expected by chance. This analysis was performed using the Partek Flow software package (34) using KEGG mouse database (35) and Fisher’s exact test is used to determine the significance of a pathway. Benjamini-Hochberg procedure (28), is applied to control the FDR. The Enrichment Score and other values were calculated to quantify the level of overrepresentation of DEGs in each pathway.

### PCA toolbox

The toolbox is designed for visualizing and analyzing PCA results. The function retrieves a correlation matrix and gene names and fills missing data points with zeros to ensure the PCA can be performed on complete data. The function calculates the cumulative variance and selects the first 25 for display on a scree plot. The function generates a 3D scatter plot of the first three principal components (PCs) by default. The visualization can be modified by using different PCs through the MATLAB code. The toolbox provides an interface for user interaction, including clustering, highlighting specific genes, and searching for genes and pathways, perform gene clustering and calculate and displays Kullback-Leibler Divergence *(KL_Div)* between the original data distribution and the PCA-resulted distribution as a measure of information loss. The details of the calculation are given in the Supplemental information.

### t-SNE toolbox

Our implemented t-distributed Stochastic Neighbor Embedding (t-SNE) toolbox that performs t-SNE calculations (36) uses the results from the correlation matrix and presents the genes based on their correlation values in 3D using a built-in ‘*tsne*’ function implemented in MATLAB. Interactive elements enable K-means clustering, visualization of brushed points, identification of individual genes and pathways, identification of close proximity genes and connecting individual genes to the STRING database.

#### t-SNE initialization

To make t-SNE analysis reproducible, the calculations were performed with the initialization step as suggested by Kobak and Berens (37) modified for 3D. The initialization process involved several steps. The data were first processed using PCA and the first three principal components (PCs) of the data were extracted. The extracted three components were standardized by dividing each by the standard deviation of the first component. This standardization ensured that the scale of the features did not disproportionately influence the results. The standardized components were then multiplied by 0.0001 which scaled down the initial positions of the data points before running t-SNE to ensure that the initial positions were close together, which insured the reproducibility of t-SNE.

#### Perplexity and other parameters

To optimize the perplexity parameter, we conducted a number of runs with varying perplexity from 5 to 200. The 3D t-SNE graphs were inspected visually to find the best cluster-like distribution of the data (**Figure S3**). For each run *KL_Div is* calculated. *KL_Div* calculates the difference between two joint probability distributions derived from the original correlation matrix (*Pm*) and from the t-SNE data (*Qm*). The method involves several steps: *i*) joint probability distribution creation, *ii*) normalization of the joint probability distribution and *iii*) *KL_div* calculations. The details of the calculation are given in the Supplemental information. *KL_div* values closer to the zero are considered to be ideal and the graph *KL_div* vs. *Perplexity* is given in **Figure S4**. Low *KL_div* value combined with the sufficient visual groupings of the scatter plot provided optimal *Perplexity* equal to 60.

Other parameters: we used *Learning Rate* = 200, that influences how the algorithm learns to map high-dimensional data to a lower-dimensional space, and the *Number of PCA Components* = 25 (since the 25 components accounted for more than 99% of all variability, see **Figure 8**).

#### Proximity mapping feature

The proximity mapping feature of the *tsne* toolbox in IVCCA provided a list of the set number of the closest genes to a searched gene name. This involved calculating the Euclidean distances between the searched gene and all other genes in the t-SNE plot, and then identified the nearest genes based on these distances.

### Venn diagram

Venn diagrams were constructed to visualize the overlap between two datasets and providing the list of the overlapped genes.

### Chromosomal distribution and histogram comparison

Each gene location on a chromosome was extracted using Partek Flow and plotted in MATLAB as histograms. The comparison between two distributions was performed using a two-sample Kolmogorov-Smirnov test (38) implemented in MATLAB.

### Network analysis

Network analysis was applied to visualize and rank genes based on their connectivity with other genes. The function implemented in the Network Analysis toolbox in IVCCA retrieved gene correlation data along with the gene names. A correlation threshold (default =0.75) was used to display the connections (edges) between the genes with pairwise correlations above the threshold level. The number of the formed connections ‘*degree*’ were calculated and made the size of the node proportional to the number of formed ‘degrees’. The Network Analysis toolbox constructed either a 2D or 3D network graph, where each node represents a gene, and edges connect genes with correlations exceeding the threshold. For better visualization, we used both node size and edge enhancement formulas Eqs. 18-19 (Supplemental Information). The equations scaled the size of each node and edge relative to the maximum degree, ensuring that nodes with more connections and higher values of edges are visually larger.

## RESULTS

### Correlation of DEGs rank genes according to their correlation values

Gene expression data from 9 mice (4 control and 5 oxaliplatin treated mice) was extracted from our earlier study (GEO repository GSE233805). Out of the 13775 identified genes in the dataset generated by RNA-seq from the hearts of mice treated with oxaliplatin and untreated controls, we selected 1744 differentially expressed genes (DEG) that satisfied the criteria: FC >1.5 and FDR <0.05, which are commonly used thresholds for determining the significance of differentially expressed genes in RNA-seq. Pearson coefficients (*q*) were calculated pairwise for each pair of genes, with the total number of unique pairwise coefficients for 1744 genes equal to 1,525,131 (see Theory, Supplementary Information). These pairwise values were plotted using the Correlation Heatmap matrix shown in **Figure 2A**. The values of *q* fall between −1 to +1, with positive values indicating positive correlation between the genes and negative values corresponding to the negative correlation. Typically, values of *q* above 0.75 or below −0.75 suggest strong correlation between the pair of genes. Interestingly, among this set of genes, more correlation coefficients between the pairs of genes were positive (see insert in **Figure 2A** showing the distribution of the pairwise correlation coefficients between the genes) suggesting that the larger number of the DEGs were positively co-regulated. A predominance of positively co-regulated DEGs might reflect underlying regulatory mechanisms that favor synergistic gene activation.

**Figure 2.**
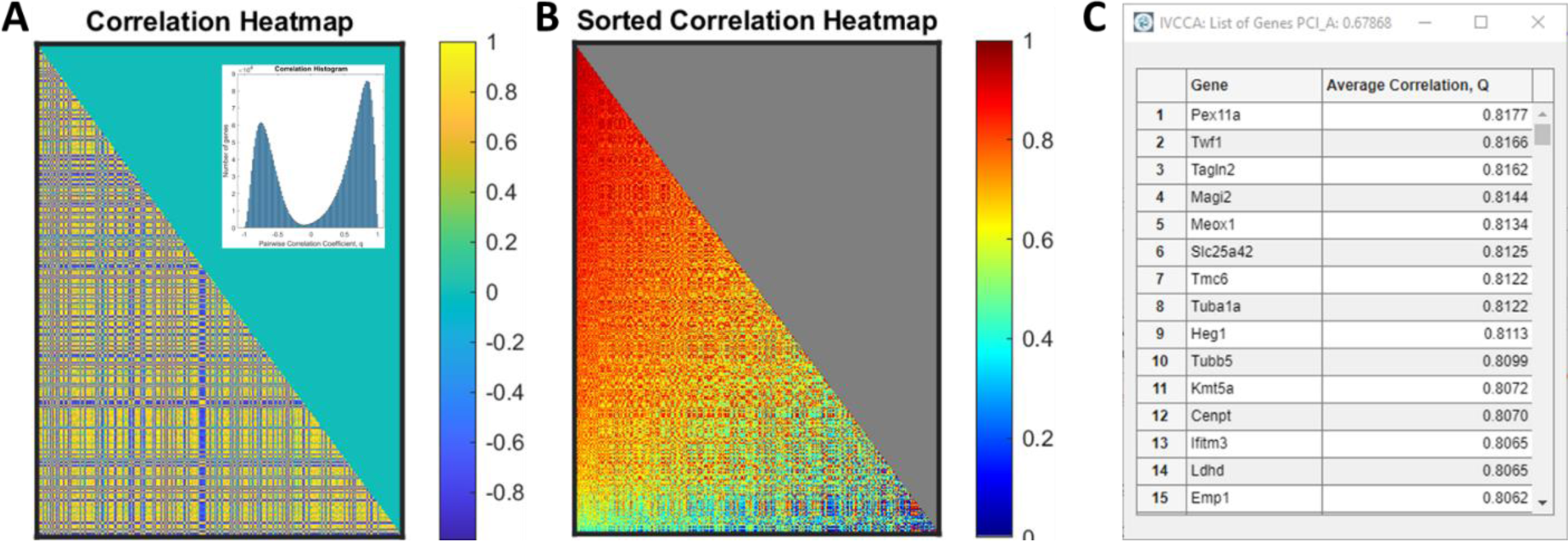
Correlation analysis of DEGs with FDR<0.05, FC>1.5 (total 1744 genes). **A**: The panel shows a correlations heatmap of genes in alphabetical order (parula colorscheme). The color reflects positive (yellow) and negative (blue) correlation between the genes. The insert shows the bimodal distribution of the gene correlation values *q* showing negative and positive correlations. The strength of correlation is defined by the colorscheme. **B**: Same map but sorted based the sum of absolute correlation values *Q*. Genes with the highest *Q* values are located in the top left corner. The strength of the correlation signal is shown by the jet colorscheme. **C**: The list of the top genes with the *Q*. *PCI_A*=0.68

The strength of correlation between individual genes can be visualized with a correlation heatmap where genes are arranged via their average global correlation coefficients (*Q*) defined as the sum of all of the absolute values of correlation coefficients divided by the total number of genes minus one to eliminate self-correlation (see Methods). The genes can then be sorted based on their *Q values* in descending order such as shown in **Figure 2B-C** highlighting the genes with the strongest associations. From this analysis, we found that more than 380 genes have values of *Q* greater than 0.75 which is typically used as a threshold for strong correlation. Such a large number of closely correlated genes might indicate a highly interconnected network of gene interactions collectively contributing to the CTX mechanism. This tightly knit network highlights the complexity of the condition and treatment, as modulating just one gene could resonate through many pathways, affecting them in largely unpredictable manner. It is this reason that we performed further pathway analysis to tease apart which genes and how those genes were modulated (below).

A similar analysis was conducted for a set of randomly selected genes without any FDR or FC thresholding (1744 genes, *RNA_seq_Random_Genes.xlsx,* Supplementary Information). The results shown in **Figure 3** are drastically different from the set of DEGs thresholded for FC>1.5 and FDR<0.05 (Figure 2). The *q* pairwise correlation values for the non-thresholded dataset were distributed more symmetrically around the zero with almost equal number of positive and negative values (**Figure 3A**, insert). The average correlation for this group of randomly selected genes was *PCI_A* = 0.37 vs 0.68 for the DEGs, suggesting lack of significant cooperation between randomly selected genes (**Figure 3B-C**).

**Figure 3.**
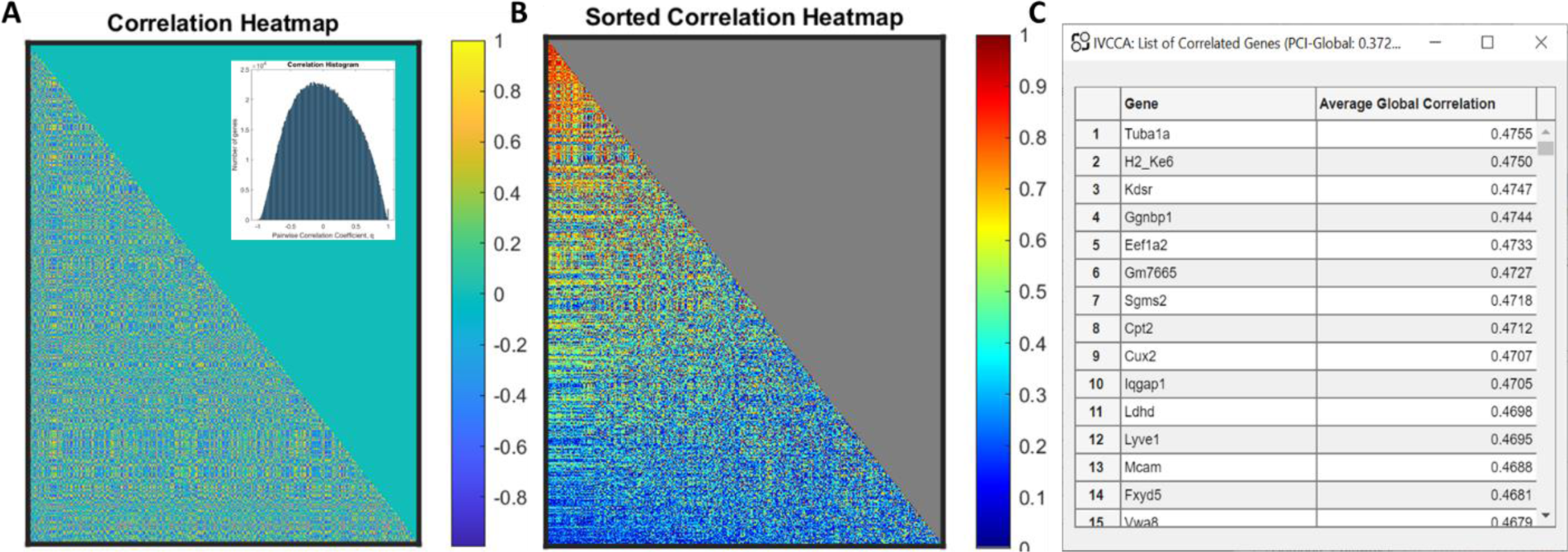
Correlation analysis of randomly selected genes (total 1664 genes). **A**: The panel shows a correlations heatmap of genes randomly generated genes (parula colorscheme). The color reflects positive (yellow) and negative correlation (blue) between the genes. The insert shows the symmetrical distribution of the gene correlation values *q* centered around zero. **B**: Correlations heatmap (jet colorscheme) sorted based on the sum of absolute correlation values *Q*. Genes with the highest *Q* values are located in the top left corner. **C**: The list of the top genes with the highest *Q* values. The average *Q* value is PCI_A=0.37

### Pathway Correlation Indices (*PCI_A* and *PCI_B*) quantifies coordinated activity of genes in a pathway

The Pathway Correlation Indices (*PCI_A* and *PCI_B*) represent the average cross-correlations of gene expressions within a specific biological pathway (*PCI_A*) and across the entire dataset (*PCI_B*). The *PCI_A* metric uses cross correlation of each gene across a specific pathway and shows how cohesive the pathway is and how coordinated the response of the genes in the pathway to the stimuli is. The *PCI_B* metric uses cross correlations of each gene across the entire dataset and reflects the position of this pathway in the hierarchy of other pathways. A large change in both metrics might suggest a coordinated response or disruption of the pathway. The PCIs range from 0 to 1, where 1 indicates a 100% correlation between the genes within the pathway or the entire dataset and 0 indicating no correlation. PCIs appears to be in general not sensitive to the FC threshold (**Figure 4**). The PCI values appear to be fairly insensitive to the FC threshold (within the same FDR threshold), although higher FC threshold strongly and expectedly decreases the number of DEGs. Low sensitivity of PCIs to the FC threshold makes this index a robust parameter to characterize and compare pathways.

**Figure 4.**
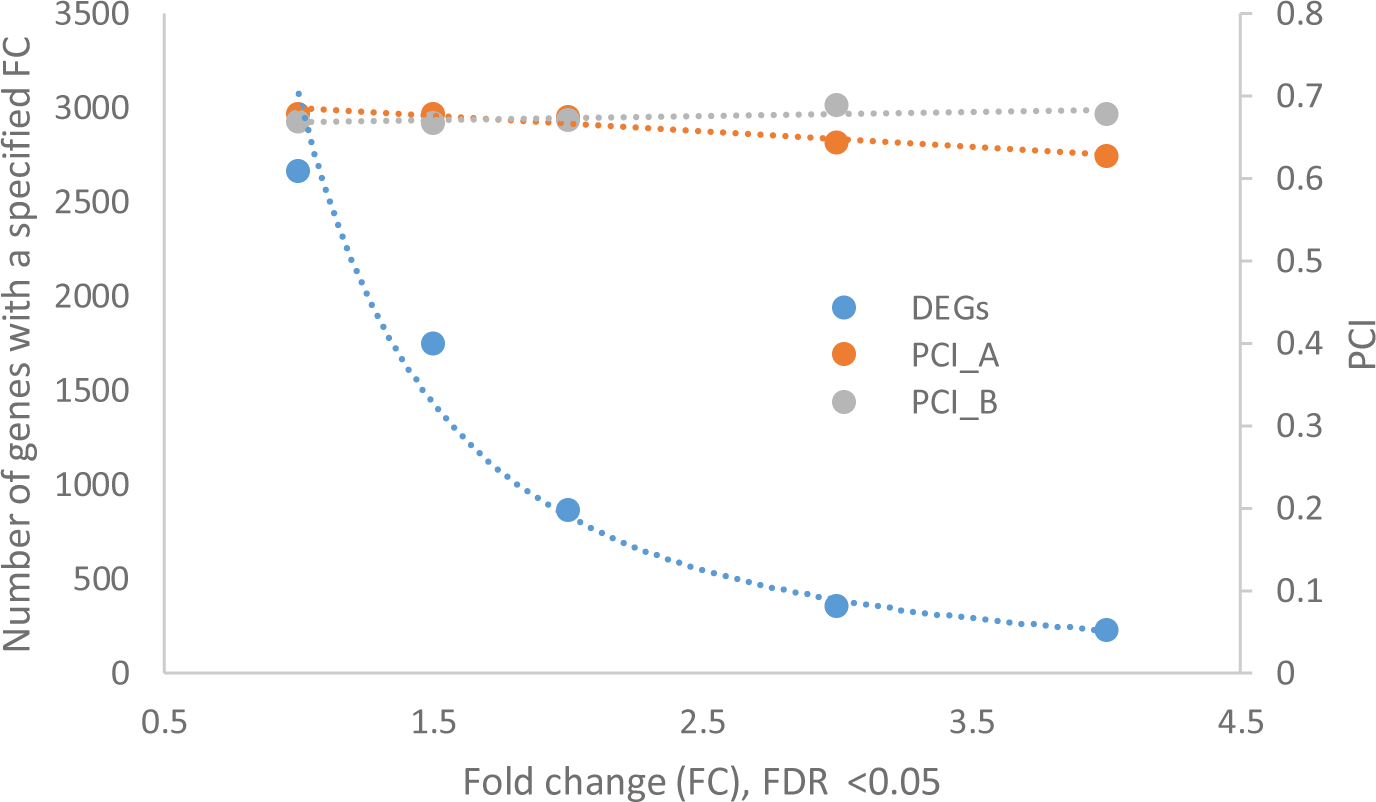
Higher FC threshold decreases the number of the identified DEGs but only weakly affects PCI_A and does not affect PCI_B. FDR<0.05 for all genes. The 13775 genes from GSE233805 dataset were selected for specific FC values (1, 1.5, 2, 3, 4) with the same FDR <0.05 threshold and then the selected genes were used in *PCI_A* and *PCI_B* calculations.

### Highly correlated DEGs are unrelated to chromosomal proximity

Genes with closely correlated values may not only imply a functional relationship but also suggest genomic proximity. Genes with high correlation might be situated near each other on a chromosome and be under the control of identical regulatory elements (39). Such co-regulated genes may have common transcription factors that bind to their promoter regions, thus enforcing their expression. To investigate whether the location of genes affect the genes distribution we mapped the 1744 DEGs (thresholded by FDR<0.05 and FC>1.5) from the mouse dataset to their respective chromosomes (data are in Supplementary Information *RNA_seq_DEGs.xlsx*). The results were compared to a set of randomly selected genes with the FDR <0.05 threshold, but no FC applied (*RNA_seq_all genes.xlsx*) and to 13775 set of identified genes in the dataset with no filtering applied (*RNA_seq_Random_Genes.xlsx*). The chromosomal distribution of DEGs mirrored the distribution of the random genes and was similar to the distribution of the entire dataset across 13,775 genes (**Figure 5**). This similarity in distributions was confirmed with a two-sample Kolmogorov-Smirnov test with the output *h* = 0 indicating that the test did not find a statistically significant difference between the two pairwise distributions [**A** (filtered genes) vs. **B** (random genes)] and [**A** (filtered genes) vs **C** (all genes)] at the 5% significance level. This result suggests that the chromosomal location of a gene does not significantly impact its likelihood of being differentially expressed by oxaliplatin and implies that a genome-wide response to oxaliplatin is not constrained to specific chromosomal regions.

**Figure 5.**
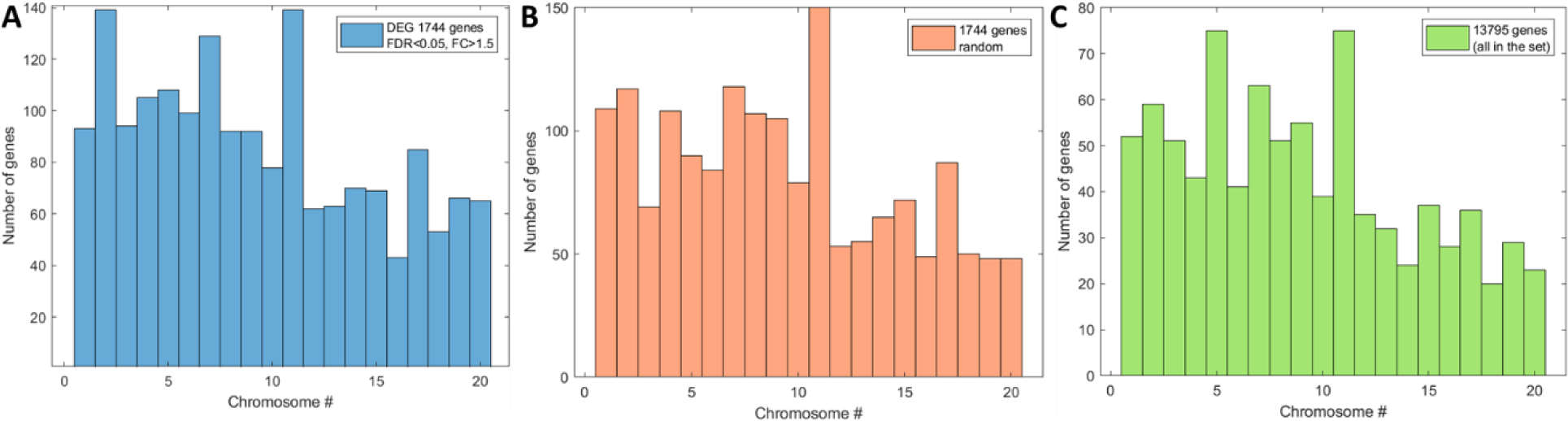
Distribution of genes on chromosome pairs. **A**: 1744 DEGs, FC >1.5, FDR<0.05, **B:** random 1744 genes and **C:** all genes (13795 genes) identified in the dataset (genes). Location on the chromosomes were assigned using Partek Flow software. Histograms were plotted with MATLAB. Pairwise Kolmogorov-Smirnov test showed *h* = 0 no statistically significant differences between the two distributions (**A** and **B**) and (**A** and **C**) at the 5% significance level. Chromosome pair #20 corresponds to X or Y.

## DISCUSSION

While the bioinformatics tools such as DAVID (40), Enrichr (41), or g:Profiler (42) as well as STRING, and COMPBIO are valuable for identifying potential targets, they have several limitations. Their effectiveness relies on the quality and coverage of the underlying databases they use. While the tools provide access to extensive databases, the completeness and timeliness of the data may vary, potentially impacting the accuracy and relevance of the results. These tools over-rely on known interactions between genes and proteins, which may limit their ability to uncover novel or uncharacterized gene targets. They do not capture the full complexity of interactions within a pathway or consider unknown genes and newly discovered genes and interactions. They are somewhat biased towards well-studied pathways, potentially overlooking important interactions or pathways relevant to new clinical conditions or preclinical models. In addition, with a few exceptions (43) these tools use average data across groups, ignoring inter-individual variability within a group. The latter is critical, since individual variations in gene expression and pathway dysregulation among members may hold important insights into potential drug targets that are not captured by these tools.

Addressing the limitation, our comprehensive interactive Inter Variability Cross-Correlation Analysis (IVCCA) tool takes into account individual expression of each differentially expressed gene. Based on the calculated correlation values between all genes in the dataset we can rank known and unknown genes and pathways, group genes into potentially new pathways and compare new pathways to existing pathway framework in a high throughput manner as discussed below. This approach ultimately enables the determination of whether a pathway or gene with potential therapeutic value can be targeted safely, without adversely affecting chemotherapy or cancer growth.

### Ranking differentially expressed genes and mapping top-ranking genes to CTX

A global correlation analysis, coupled with ranking the genes according to their PCI scores, identified the top 1% genes (top 20 genes): *Pex11a, Twf1, Tagln2, Magi2, Meox1, Slc25a42, Tmc6, Tuba1a, Heg1, Tubb5, Kmt5a, Cenpt, Ifitm3, Ldhd, Emp1, H2-Ke6, Tspan2, Extl1, Mospd2, Arpc1b*) (**Figure 2**C). These genes have a diverse functional range. Key functions include regulation of vasculature (*Heg1, Tagln2*), metabolism (*Pex11a, Slc25a42, Ldhd*), cell structure and integrity (*Twf1, Tagln2, Tuba1a, Tubb5, Meox1*), cell cycle and chromatin organization (*Tuba1a, Tubb5, Kmt5a, Cenpt*), and immune response (*Ifitm3*). The expression profiles of some genes are shown in **Figure S5**.

Many of these top-ranking genes are well recognized for their involvement in heart. For example, *Heg1* (FC=1.89), a key protein in endothelial cell biology, is notable for its role in cardiovascular development and function (44). Alteration in the HEG1 protein has been linked to vascular abnormalities in zebrafish and mice (44,45). *Tagln2*, another top-ranking gene, encodes a protein that regulates smooth muscle cell function. This gene is significantly overexpressed (FC=2.69) in the heart tissue of oxaliplatin treated animals. A similar pattern of overexpression has been also seen with another chemotherapy drug doxorubicin (46). *Pex11a*, which is moderately expressed in heart tissue, is a vital protein involved in linking peroxisomal membranes to motor proteins necessary for perixosomal replication (47). In mice treated with oxaliplatin, *Pex11a* is downregulated (FC= −1.88), that aligns with the general downregulation of the fatty acid (FA) metabolism pathway in the heart as we previously reported (30). Another key gene, *Slc25a42*, encodes a protein responsible for importing coenzyme A (CoA) into the mitochondrial matrix (48). This process is crucial for energy metabolism in heart cells, as CoA is essential for various metabolic processes, including FA synthesis and oxidation. The downregulation of *Slc25a42* (FC=-1.79) suggests a reduced CoA level in mitochondria, potentially leading to lactic acidosis and myocardial weakness (48,49). Similarly, *Ldhd*, a gene encoding mitochondrial lactate dehydrogenase D involved in D-lactate metabolism (50), is also downregulated (FC= −2.20). Since *Ldhd* is involved in D-lactate processing instead of L-lactate (48), this decrease in expression suggests impaired D-lactate processing, leading to harmful D-lactate accumulation in the heart. Notably, all three genes — *Pex11a*, *Slc25a42*, and *Ldhd* — show a high degree of correlation, with pairwise correlation values exceeding 0.99, suggesting high level of coordination between these genes.

Several top-ranking genes are involved in maintaining the structural integrity of cells and tissues, with many of these genes being overexpressed. Their change in expression is often seen in damaged heart tissues. For instance, *Twf1*, (FC=1.68) that encodes an actin monomer-binding protein has also been observed in hypertrophic myocytes *Meox1*, crucial for muscle development, also exhibits significant upregulation (FC=3.41). Increases in its expression is often observed in mouse models and patients with hypertrophic cardiomyopathy (51) and used as a target for treating cardiac fibrosis (52).

Oxaliplatin primarily works by forming platinum-DNA adducts. These adducts cause DNA cross-linking, which interferes with DNA replication and transcription, ultimately disrupting the cell cycle. It is not surprisingly that several high-ranking correlation genes are part of the cell cycle and cell division pathways including senescence, p53, apoptosis, and others. In response to stress, cells initiate various protective mechanisms, including the upregulation of certain pathways that promote survival, which may involve genes associated with cell division. Cardiomyocytes and other heart tissue forming cells are in generally postmitotic. After the early stages of life, the mammalian heart has a very limited capacity to generate new cardiomyocytes (53). Instead, the heart myocardium responds to injury by remodeling rather than regeneration through cell division. Increases in mitotic activity of endothelia cells in the oxaliplatin treated mice are associated with the spindle which are constructed from α-tubulin (*Tuba1a*) and β-tubulin (*Tubb5)* dimers (54). *Tuba1a* produces an alpha-tubulin monomer. *Tubb* (T*ubb5*) produces a beta-tubulin monomer. The spindle fibers of the mitotic spindle apparatus are formed from α- and β-tubulin dimers. The elevation of *Tuba1a* and *Tubb5* are most likely due to mitotic activity in this case in the endothelia since that tissue gets hit by oxaliplatin due to the high turnover and endothelia, unlike cardiomyocytes, are capable of replication.

Another high-ranking gene *Cenpt* (FC=2.41) is part of the centromere protein family (CENP) that are involved in chromosome segregation during cell division. CENP relationship with tubulins is well known (55,56), as centromeres, where CENP proteins are located, attach to spindle fibers composed of microtubules. Our Gene-to-Pathway analysis show that both types of genes are synchronized in their response to oxaliplatin with a high average correlation above 0.8 (**Figure S6**). Another gene that is involved in the regulation of chromatin is *Kmt5a* (FC=-1.90) that regulates lysine-20 methylation in histone H4. Recent results showed a direct link between the lysine methylation at this site and the heart failure in mice and humans (57).

In summary, the top-ranking genes with high average correlation indices point to several diverse pathways influenced by oxaliplatin. These pathways evidence both the body responses to CTX and the resultant remodeling of heart tissue. This remodeling, involving processes like fibrosis and hypertrophy, could contribute to the observed progression of heart failure.

### Ranking pathways based on Pathway Correlation Indices

Finding the most relevant pathways that can describe biological processes in the heart of the oxaliplatin treated mice is one of the key challenges to understand the mechanism of CTX. We address this challenge by calculating Pathway Correlation Indices (*PCI_A* and *PCI_B*) defined as the average of absolute individual pairwise correlations for all genes within the pathway (*PCI_A*) or average of the absolute value of the correlations for genes in the pathway with the genes in the entire dataset (*PCI_B*) (Eq.5, Supplemental Information). **Figure S2** illustrates the difference between the two metrics. Pathways with the high *PCI* values exhibit the strongest coordinated gene expression response to oxaliplatin pointing to pathways that are especially active under the oxaliplatin treatment. We also define PAI that quantifies the extent to which a particular pathway is activated and is defined as the ratio of differentially expressed genes found in the set to the total number of genes in the pathway. The *Z-score* accounts for both *PCI_B* and *PAI* (Eqs. 14-16, Supplemental Information). Our GUI-implemented algorithm enables screening hundreds of pathways and calculate their Z-scores within seconds.

The top IVCCA results of the screening of more than 340 pathways from the KEGG database and 15 relevant literature-based pathways is shown in **Figure 6** with the few significant tabulated in **Table 2**. The top-ranking pathway from the screens was the Metabolic Energy pathway that we have defined in (30) with the highest Z-score. This is in line with what we and others have reported for chemotherapy drugs in mice and humans (30,58). Metabolic changes from chemotherapies lead to weight loss, fatigue, appetite changes, and may have long-term effects on cardiac function depending on the severity of the metabolic changes (59,60).

**Figure 6.**
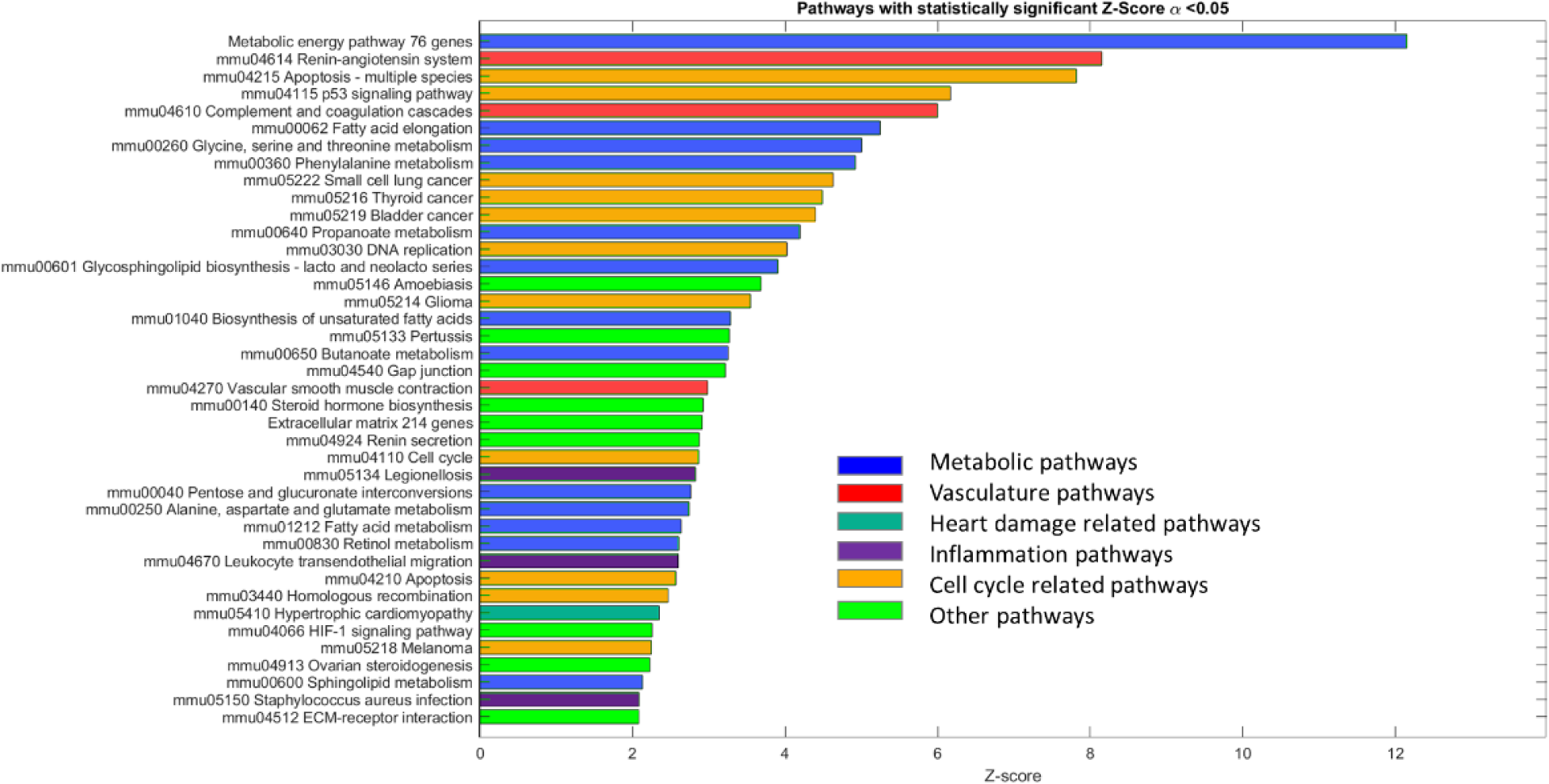
IVVCA ranking of KEGG (344 pathways) and literature-based pathways based on the Z-score. Only statistically significant pathways above the Z-score critical are shown. Colors mark pathways related to the same group.

Among the prominent pathways identified were the renin-angiotensin system (RAS), apoptosis, and the p53 signaling pathway. It was not surprising to see apoptosis and the p53 pathway ranked highly, as they are well-known to be involved in cell division processes affected by oxaliplatin. However, the identification of RAS as a top-ranking pathway was unexpected but significant. RAS is fundamentally important in regulating blood pressure and renal sodium retention and plays a critical role on the health and disease of the cardiovascular system. Renin, an enzyme produced by the kidneys in response to poor nephron filtration due to low blood perfusion, converts plasma angiotensinogen (*Agt*) into angiotensin I which is further converted by the angiotensin I, converting enzyme (*Ace*) into angiotensin II by the angiotensin II converting enzyme (*Ace2*). All three genes in this pathway were upregulated, *Agt* (FC=3.20), *Ace* (FC=1.71) and *Ace2* (FC=3.28) (61). Chronic elevated levels of angiotensin II leads to hypertrophy of the heart muscle and fibrosis in mice (62). Because of its role in heart injury, the RAS is a target for several heart medications where a variety of ACE inhibitors and angiotensin receptor blockers are commonly used to reduce the risk of heart damage (63). Given the intrinsic connection between the RAS in linking kidney function and heart health, it is not surprising that oxaliplatin is associated with nephropathy, a recognized condition in patients undergoing treatment with this drug (64,65).

It is interesting to compare the top pathways found by IVCCA with the top pathways founds by the existing methods. The KEGG enrichment analysis implemented in the Partek Flow software analysis identified 22 statistically significant pathways (20 from KEGG and 2 from custom) with top-ranking pathways that include vasculature regulation, inflammation, and cancer related pathways (**Figure S7)**. Other significant pathways included neuroactive ligand-receptor interaction, RAS, and metabolic pathways. Out of these 20 KEGG pathways there were 9 that were also identified by IVCCA suggesting that despite the difference in methodology both strategies lead to similar results. While the overlapping results speak to the validity of our approach, our new tool has the potential to recognize previously unknown pathways.

The correlation analysis of the pathways from the GO database were consistent with KEGG and gave a similar pathway ranking landscape (**Figures S8-S10** show ranking pathways from GO biological functions, GO cell components, GO molecular functions databases based on their Z-scores). From the screening of more than 1700 GO pathways that contain more than 5 genes in that particular pathway, IVCCA identified 167 statistically significant GO pathways. Similar to KEGG, the top-ranking GO biological functions identified by IVCCA were related to cell division, regulation of cardiac activity, and actin filament organization. Others also included blood vessel-related pathways, regulation of the metabolic processes, and inflammation.

### Finding new pathways and target genes

Investigating a relatively unexplored physiological condition such as CTX introduces challenges in understanding its underlying mechanisms, as the existing established pathways (KEGG, GO and others) may not be sufficient to explain the observed physiology. The unique characteristics of oxaliplatin induced CTX requires exploring novel pathways that have not been traditionally associated with similar physiological processes. We approached this challenge through clustering of the DEGs based on their correlation values. We implemented three methods to identify the clusters: dendrogram, principal component analysis (PCA) and t-Distributed Stochastic Neighbor Embedding (t-SNE). The results contrasting the advantages and disadvantageous of these methods are discussed below.

#### Dendrogram produces clusters that are either too small or too large

Within a dendrogram built from correlation data, gene clusters located closely together exhibit greater similarity in terms of their correlation patterns compared to clusters positioned farther apart. By adjusting the threshold level (i.e., distance), the number of clusters can be defined, and color-coded for better visualization (**Figure 7**). One of the challenges in this process is to find the number of clusters that faithfully represent the data. This task is subjective and usually comes from the visual inspection of the dendrogram. More objective techniques to identify the number of clusters is to use algorithms such as an elbow (31) and a silhouette (32) to visual those relationships. The elbow technique plots the variance as a function of the number of clusters with the “elbow” of the curve represents an optimal number for clustering. Silhouette technique plots how close each point in one cluster is to the points in the neighboring clusters with a high value indicates that the object is well matched to its own cluster and poorly matched to neighboring clusters. In the given case with 1744 DEGs dataset, the elbow method did not show the obvious ‘elbow’ point (**Figure 7A)**, while the silhouette method suggested either 6 or 10-15 clusters to be optimal (**Figure 7B**). Based on this suggestion, the dendrogram was thresholded with the Distance = 8.5 resulting in 10 clusters (**Figure 7C**).

**Figure 7.**
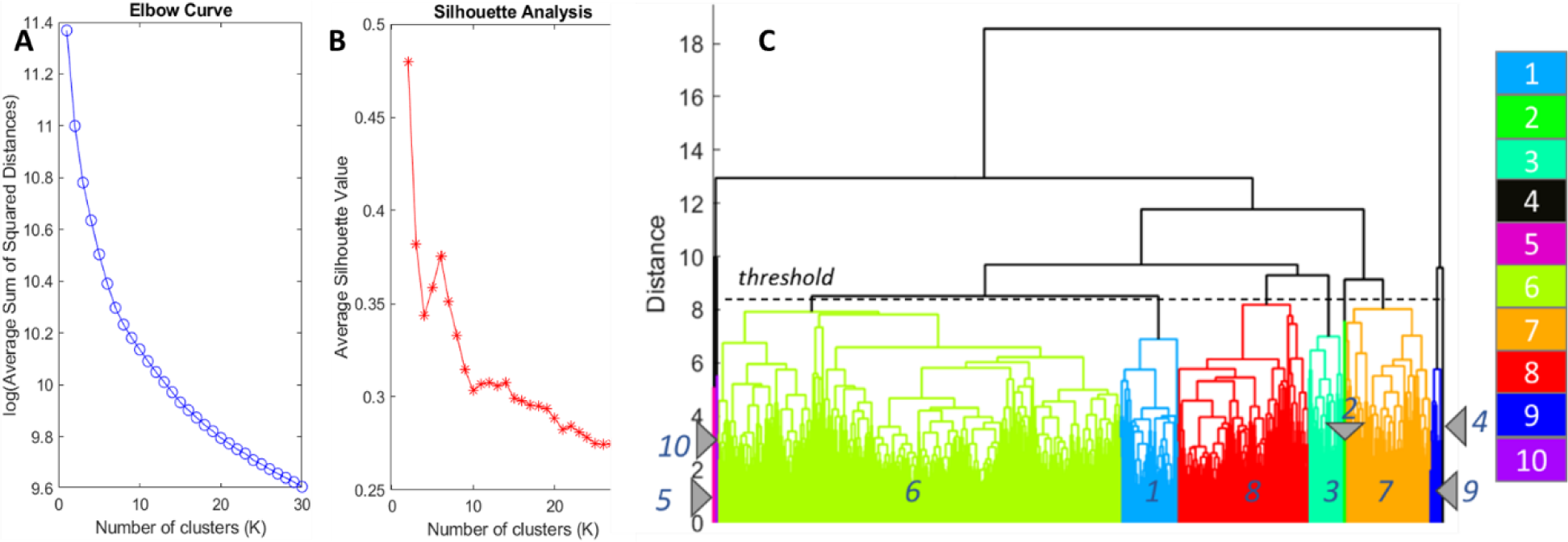
Clustering with dendrogram. Elbow (**A**) and silhouette (**B**) methods were used find the optimum number of clusters. The curves were calculated 5 time and averaged. Elbow method was unable to identity the number of clusters. Silhouette method suggests 6 or 10-15 clusters. **C:** Dendrogram of the genes based on their correlation values. The number of unique clusters were defined by the threshold distance =8.5 resulting in 10 identified clusters. Genes with higher correlation to each other belong to the same cluster. The clusters sizes are highly different ranging from 1 to 916 genes per cluster. Genes from the 1744 DEGs filtered for FDR<0.05 and FC>1.5.

As seen in **Figure 7C**, the resulting dendrogram produced clusters ranging from very large (above 900 genes) to very small (1-2 genes). Ideally, clusters should contain 50 to 200 genes for conventional databases such as STRING that covers KEGG, GO and other databases. Large clusters (such as Cluster #6 with 961 genes) risk incorporating noise or false positives. Small clusters with only few genes (such as clusters #2, 4, 5, 9, 10) fail to show significant enrichment. Therefore, only a few clusters gave meaningful results. Cluster #1 (138 genes in the cluster) was linked to the pathways primarily related to the regulation of the cell cycle. Cluster #7 showed mixed pathways including FA metabolism, angiogenesis, and complement activation. Cluster #8 also represented a mix of pathways such as regulation of blood vessels and response to hyperoxia among others.

#### PCA does not produce clusterisable pattern

Principal Component Analysis (PCA) of the correlation matrix provides a different approach to visualize the overall structure of gene-to-gene relationships and identify clusters. In this process, the principal components (PCs) represent the directions of maximum variance in the data, essentially condensing the information into fewer dimensions while retaining the relationships between the genes. This typically enables the identification of patterns and clusters among genes. The PCA analysis of the correlation matrix with 1744 genes provides 1744 PCs. The graph of the cumulative variance vs first 25 PCs is shown in **Figure 8**A showing that 85% variability lies within the first three PCs and 99.9% for the first 25 PCs. The PCA scatter plot of the first three PCs is shown in **Figure 8B**, where each point represents a particular gene, and its location corresponds to correlation with all other genes aligns along these PCs.

**Figure 8.**
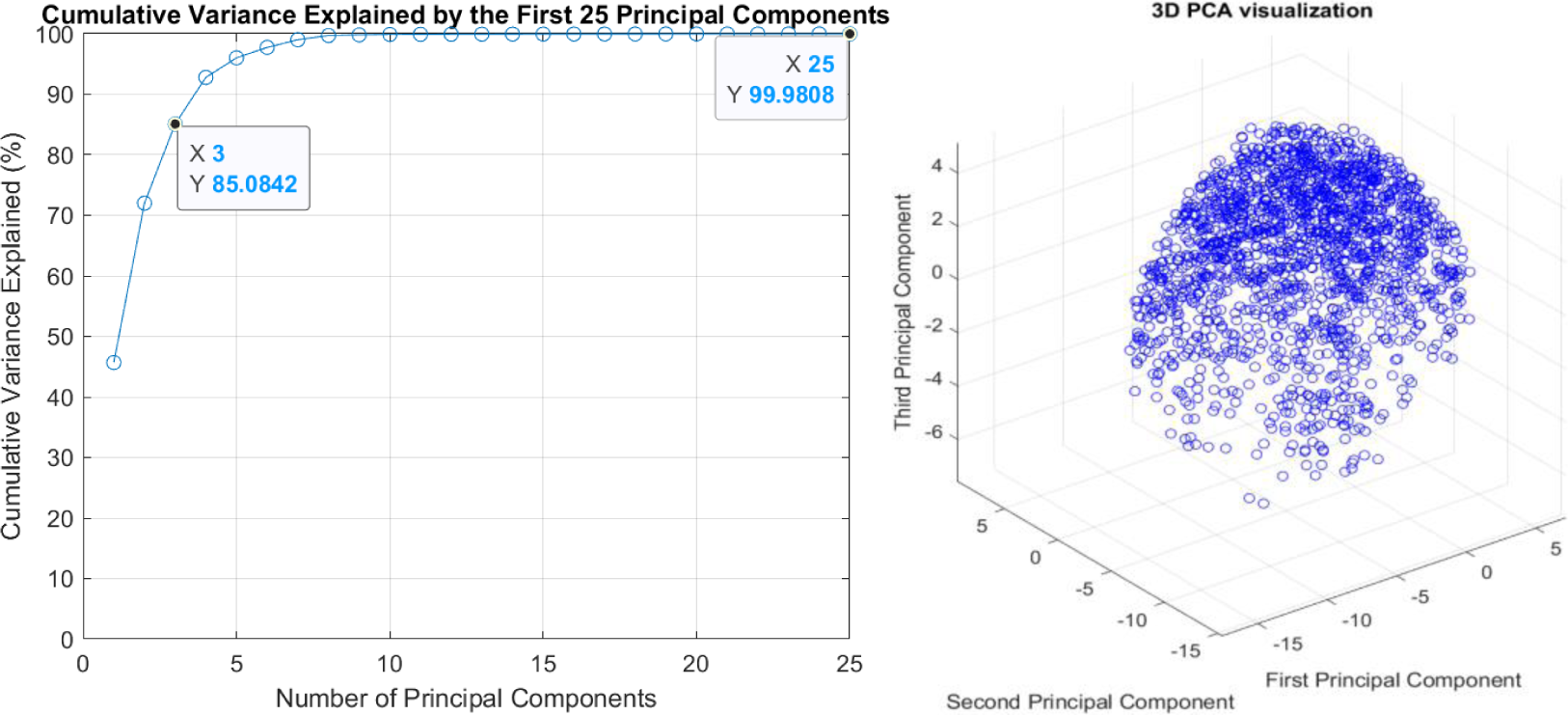
PCA of the correlation matrix composed of 1744 DEGs. **Left**: Cumulative variance plot (percent) for the first 25 PCs. **Right:** PCA visualization for the first three PCs in a 3D scatter plot of the 1744 DEGs.

The PCA plot with the first three PCs in a 3D plot makes the data to appear like a ball without much discernible structure. A close to spherical distribution might indicate that there are no strong patterns or clusters captured by these components. The scatter plots with higher-level PCs were similar with no clear groupings (not shown). The lack of apparent structure makes reasonable clustering of the PCA plot difficult and prompted us to search for different clustering technique.

#### t-SNE provides a solution to cluster correlation data to identify CTX related clusters

To achieve better clustering, we applied t-SNE, a well-known machine learning algorithm designed for visualizing high-dimensional data in a lower-dimensional space (36). This non-linear technique is particularly suitable at capturing non-linear relationships between genes, which PCA might miss since PCA is a linear method. Being a stochastic method t-SNE starts randomly, leading to varying results affecting the locations of the genes in the visualization plot. To minimize the randomness, we followed a modified Kobak and Berens suggestion (37) by initiating t-SNE calculations from the first PCs. Since we used a 3D representation, we started from the first three PCs from the PCA calculations. Beginning with PCA as the initialization made t-SNE less sensitive to random initializations, leading to more consistent and reproducible results. This approach allowed us to optimize perplexity, a crucial tuning parameter that significantly affects data point positioning in scatter plots and, consequently, the identification of clusters or pathways. We varied perplexity from 5 to 200 by evaluating outcomes visually and quantitatively using the Kullback-Leibler Divergence (*KL_div*) score. Lower *KL_div* scores indicate better embeddings. Our analysis, including visual observations of 3D scatter plots (shown in **Figure S3**) and *KL_div* values (**Figure S4**) determined that the optimal perplexity value was 60. At this level, the data points showed clear patterns and significant clustering (**Figure 9**).

**Figure 9.**
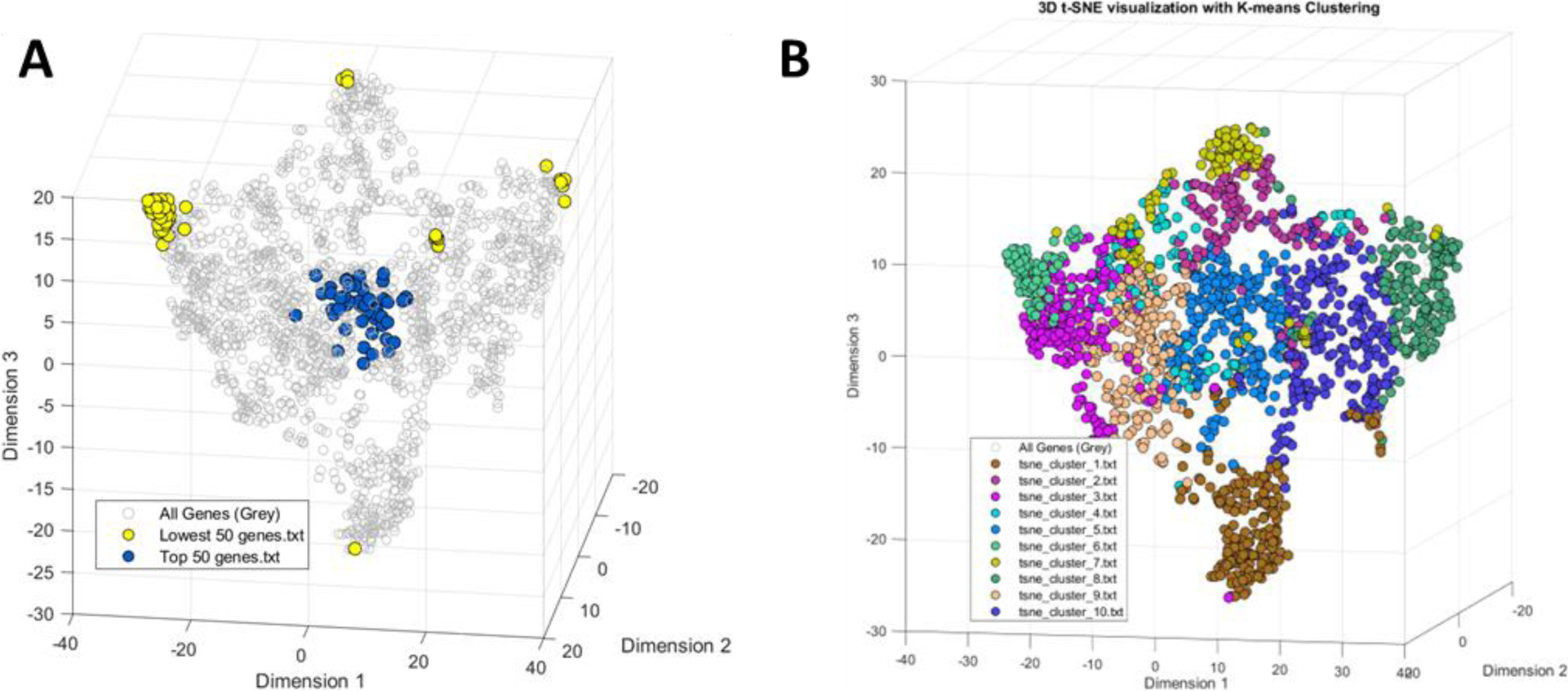
t-SNE results of 1744 DEGs. **A:** the distribution of 50 genes with the highest and lowest average correlation values. **B:** the distribution of 10 clusters as defined by K-means. Major optimized parameters: perplexity = 60, learning rate = 200.

In the resulting t-SNE 3D scatter plots, closely positioned points indicate strongly correlated data, while distant points suggest weaker correlations. For instance, the top-ranking 50 genes with the highest average correlation values (*Q*) form a distinct cluster in the t-SNE plot (**Figure 9A**), contrasting with the multiple widely dispersed clusters formed by the 50 genes with the lowest *Q* values. Notably, clusters corresponding to some known pathways from KEGG, GO, and others are discernible on the t-SNE plot (Figure S1C or **Figure 9B**). Pathways related to the extracellular matrix, centromeric regions, and energy metabolism all show distinct localizations (**Figure S11**), suggesting t-SNE’s potential in identifying novel pathways based on their placement in the 3D scatter plot.

K-means clustering of the t-SNE scatter plots enables the identification of inherent groupings within the data. As with the dendrogram and PCA methods, we generated ten clusters. The clustering of the t-SNE data yielded more uniform clusters containing reasonable number of genes between 87 to 240 genes (**Figure 9B)**. The genes in each cluster were analyzed whether they correspond to known KEGG or GO pathways using STRING database. The results of this analysis with the Z-score (Eqs. 16-17, Supplemental Information) for each cluster is shown in **Table 1**.

**Table 1.** K-means clustering of the t-SNE 3D plot and STRING analysis of the clusters. Column cluster # corresponds to the cluster on the t-SNE map. STRING assignment column shows the numbers of significantly enrichment clusters by GO, KEGG with brief names of the top pathways (STRING interaction score >0.4).

| Cluster # | Number of genes in cluster | # GO pathways | # KEGG pathways | STRING assignment | Z-score |
| --- | --- | --- | --- | --- | --- |
| 1 | 183 | 192 | 5 | DNA replication, cell cycle related pathways | 29.7 |
| 2 | 133 | 38 | 26 | Immune response, inflammation, response to hypoxia, regulation of heart rate | 29.1 |
| 3 | 220 | 22 | 1 | Complement and coagulation cascade, collagen metabolism, extracellular matrix | 26.8 |
| 4 | 121 | 2 | 0 | Protein phosphorylation | 24.4 |
| 5 | 249 | 47 | 0 | Actin cytoskeleton, fiber organization | 27.7 |
| 6 | 87 | 3 | 1 | Infection, complement cascade | 21.0 |
| 7 | 110 | 1 | 0 | Aging, myosin filament | 30.9 |
| 8 | 169 | 47 | 0 | Apoptosis, response to hyperoxia | 28.1 |
| 9 | 227 | 15 | 0 | Muscle cell development, vascular system | 26.7 |
| 10 | 245 | 105 | 3 | Regulation of muscle contraction | 30.7 |

The Z-score ranking of the clusters indicated their relative representation in the CTX. All clusters were statistically significant, exhibiting higher Z-scores primarily because all genes in the clusters are DEGs. Some clusters indicated relationship to the known cardiac related pathologies, for example clusters #2, and #9 were directly associated with heart-related issues, containing a larger number of genes known to correlate with heart conditions.

### Avoiding chemotherapy targeted and cancer pathways with Pathway-to-Pathway correlation analysis

A key objective in managing chemotherapy-induced CTX is to mitigate heart damage without interfering with the chemotherapy effectiveness in eradicating cancer and. This challenge brings a complex triangle among the treatment of the tumor, minimizing CTX, and the chemotherapy agent. To this end, we implemented a Pathway-to-Pathway comparison approach using a cosine similarity (CS) score as a metric to quantify the relationship between the reference pathways and the pathways of interest. Two pathways were used as references and all other pathways were compared against them. Given that chemotherapy agents are frequently designed to target the cell cycle to stop cancer cells from multiplying, we used the KEGG mmu04110 cell cycle pathway as the first reference pathway. Since oxaliplatin is a first line chemotherapy agent for the treatment of colorectal cancer, we used the colorectal cancer pathway (KEGG mmu05210 colorectal cancer) as the second reference.

Pathways exhibiting an extensive gene overlap or significant gene-to-gene pairwise correlations between the pathway and the reference yield higher CS scores, with a maximum of CS = 1 indicating complete overlap. We envision, that the genes in these pathways should be avoided when minimizing the effect of chemotherapy on CTX. Weakly correlated pathways show low CS scores. The genes in those low CS pathways might present a significant interest as they are expected to be relatively safe to use. The algorithms implemented in IVCCA allows the researcher to make the comparison screening of hundreds of pathways quickly against a single reference pathway. The results for the selected pathways versus the cell cycle (mmu04110) or versus colorectal cancer (mmu05210) reference pathways from the KEGG, literature-based pathways as well from t-SNE clusters are summarized in **Table 2**.

**Table 2.** Pathway comparison to mmu04110 Cell cycle and mmu05210 Colorectal cancer using cosine similarity. Higher values correspond to higher similarity. Color labeling: orange colors represent high score CS>0.6 in any of the reference comparison indicating strong interference with any of the two references; blue color indicates moderate interference 0.5<CS<0.6, green color indicates low level of interference with CS<0.5. Data are presented in CS value/number of overlapped genes between two selected pathways.

| Potential therapeutic pathway | Cosine Similarity Score (CS) to mmu04110 Cell cycle pathway | Cosine Similarity Score (CS) to mmu05210 Colorectal cancer pathway |
| --- | --- | --- |
| mmu04110 Cell cycle | 1.00/125 | 0.569/5 |
| mmu05210 Colorectal cancer | 0.569/5 | 1.00/85 |
| **Heart related** |  |  |
| mmu04260_Cardiac muscle contraction | 0.479/0 | 0.535/0 |
| mmu04261_Adrenergic signaling in cardiomyocytes | 0.494/0 | 0.533/0 |
| mmu04614_Renin-angiotensin system | 0.533/0 | 0.503/0 |
| Arrhythmia inherited (66) | 0.330/0 | 0.488/0 |
| SAN pacemaker (67) | 0.388/0 | 0.488/0 |
| mmu05414_Dilated cardiomyopathy | 0.458/0 | 0.477/0 |
| Cardiac hypertrophy_26 genes (68) | 0.375/0 | 0.430/0 |
| Cardiac myopathy_18_genes (69) | 0.345/0 | 0.418/0 |
| mmu05414_Dilated cardiomyopathy | 0.453/0 | 0.478/0 |
| mmu04020_Calcium signaling pathway | 0.508/0 | 0.530/1 |
| mmu05412_Arrhythmogenic right ventricular cardiomyopathy | 0.442/0 | 0.484/0 |
| **Blood vessel related** |  |  |
| mmu05417_Lipid and atherosclerosis | 0.549/1 | 0.405/0 |
| mmu04270_Vascular smooth muscle contraction | 0.491/0 | 0.518/0 |
| mmu04610_Complement and coagulation cascades | 0.442/0 | 0.405/0 |
| **Metabolism related pathways** |  |  |
| mmu00604_Glycosphingolipid biosynthesis - ganglio series | 0.650/0 | 0.709/0 |
| mmu00061_Fatty acid biosynthesis | 0.707/0 | 0.703/0 |
| mmu00620_Pyruvate metabolism | 0.631/0 | 0.688/0 |
| mmu00010_Glycolysis Gluconeogenesis | 0.570/0 | 0.570/0 |
| Metabolic Energy (30) | 0.501/0 | 0.538/0 |
| mmu01212_Fatty acid metabolism | 0.516/0 | 0.515/0 |
| mmu00062_Fatty acid elongation | 0.485/0 | 0.507/0 |
| mmu00760_Nicotinate and nicotinamide metabolism | 0.459/0 | 0.471/0 |
| **Inflammatory pathways** |  |  |
| mmu04668_TNF signaling pathway | 0.495/0 | 0.525/2 |
| mmu04660_T cell receptor signaling pathway | 0.488/0 | 0.550/1 |
| mmu04217_Necroptosis | 0.548/0 | 0.546/1 |
| Inflammation (70) | 0.473/0 | 0.521/1 |
| mmu04670_Leukocyte transendothelial migration | 0.537/0 | 0.541/0 |
| mmu04062_Chemokine signaling pathway | 0.551/0 | 0.526/0 |
| **Other relevant pathways** |  |  |
| Transcription factors (71) | 0.512/3 | 0.520/3 |
| mmu04530_Tight junction | 0.535/1 | 0.561/2 |
| Epigenetic (MGI search) | 0.540/3 | 0.497/2 |
| mmu04066_HIF-1 signaling pathway | 0.496/3 | 0.544/1 |
| mmu04211_Longevity regulating pathway | 0.500/1 | 0.522/2 |
| mmu04540_Gap junction | 0.589/1 | 0.600/1 |
| mmu04810_Regulation of actin cytoskeleton | 0.555/0 | 0.556/1 |
| mmu04020_Calcium signaling pathway | 0.503/0 | 0.530/1 |
| mmu04714_Thermogenesis | 0.584/0 | 0.586/0 |
| mmu04146_Peroxisome | 0.541/0 | 0.592/0 |
| mmu04710_Circadian rhythm | 0.538/0 | 0.636/0 |
| Extracellular matrix (29) | 0.446/0 | 0.410/0 |
| **Cell division and survival** |  |  |
| mmu04218_Cellular senescence | 0.623/8 | 0.588/5 |
| mmu03410_Base excision repair | 0.728/0 | 0.633/0 |
| mmu04115_p53 signaling pathway | 0.642/8 | 0.604/7 |
| mmu03030_DNA replication | 0.703/3 | 0.602/0 |
| mmu04210_Apoptosis | 0.543/2 | 0.574/8 |
| **t-SNE clusters** |  |  |
| tsne_cluster_1.txt | 0.682/12 | 0.511/2 |
| tsne_cluster_2.txt | 0.508/0 | 0.574/2 |
| tsne_cluster_3.txt | 0.362/0 | 0.337/0 |
| tsne_cluster_4.txt | 0.412/1 | 0.477/1 |
| tsne_cluster_5.txt | 0.656/3 | 0.649/1 |
| tsne_cluster_6.txt | 0.141/0 | 0.165/0 |
| tsne_cluster_7.txt | 0.308/1 | 0.367/1 |
| tsne_cluster_8.txt | 0.451/1 | 0.547/3 |
| tsne_cluster_9.txt | 0.559/2 | 0.515/1 |
| tsne_cluster_10.txt | 0.644/5 | 0.646/2 |

The range of the calculated CS scores were from 0.155 (mmu05033 Nicotine addiction) to 0.730 (mmu03410 Base excision repair). Based on the results we grouped the pathways in three groups based on their SC values: above 0.6, between 0.5 and 0.6 and below 0.5. Pathways with SC above 0.6 have strong overlap with any of the reference pathways and should be avoided, pathways with CS between 0.5 and 0.6 are moderately risky, and below 0.5 might be relatively safe to target. As expected, pathways related to the cell cycle such senescence, p53 and DNA repair showed high level of CS>0.6 for both references with a significant gene overlap. Similarly, several t-SNE defined clusters such as #1, #5, and #10 also show strong correlation with the reference pathways. The high degree of similarity with the reference pathways suggests that interventions in the gene(s) in these pathways could inadvertently suppress oxaliplatin therapeutic effect as well as potentially promote cancer growth. Most heart-specific pathways are close or fall below CS = 0.5 threshold, allowing for safe treatment of cardiotoxicity to improve cardiac performance. Only a few metabolic pathways are also relatively safe to target. Specifically, the treatment of the nicotinate and nicotinamide metabolism might be especially beneficial to the CTX patients without the risk of decreasing the efficiency of chemotherapy or promoting cancer growth. Similarly, targeting pathways described by t-SNE clusters #3, 4, 6 and 7 seems safe to apply. Future experiments in cancer models in animal model systems will examine whether this line of treatment will minimize CTX during chemotherapy without aiding the cancer.

#### Identifying target genes through network analysis

Following the results of the Pathway-to-Pathway correlation the next step is to identify the suitable target genes in these pathways. These genes can be identified visually from the STRING database, or through methods implemented in Weighted Gene Co-expression Network Analysis (WGCNA) (72). We rationalize that similar information can be extracted using the IVCCA approach using network analysis visualization tool.

In this network-based visualization, genes are represented as points (nodes) and their correlations as lines (edges) connecting them (**Figure 10A**). Highly connected genes (nodes) with a high number of interactions (a high degree node) above a certain threshold (*Q* here defined as 0.75) can be considered as hub genes. As an example, we performed the network analysis for the cardiac Metabolic Energy Pathway (30). This pathway currently comprised of 76 genes involved in FA oxidation, glycolysis, amino acid metabolism and NAD synthesis in the hearts of oxaliplatin treated mice. Filtering these genes for FDR<0.05 and FC>1.5, the number of genes is reduced to 36 DEGs.

**Figure 10.**
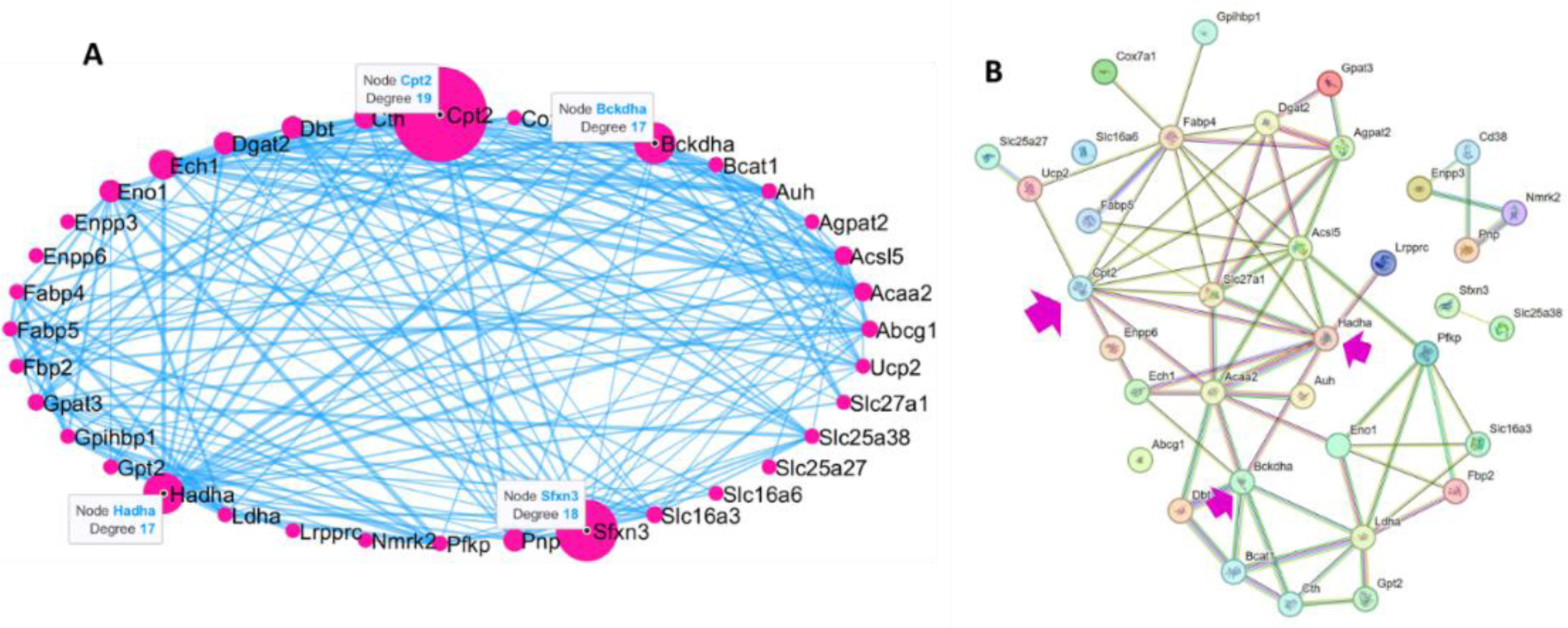
Network analysis of 36 DEGs from the energy metabolism pathway. **A (IVCCA network analysis)**: The thickness of the edges (lines) corresponds to the absolute correlation value between the connected genes. The size of the nodes represents the number of connections to other genes with an absolute correlation above the threshold of 0.75. Among the top genes identified in this analysis are *Cpt2* (19 connections), *Sfxn3* (18 connections), *Bckdha* (17 connections) and *Hadha* (17 connections). **B (STRING network analysis)**: The top genes are defined by connectivity, measured by node degree (number of connections). The threshold interaction score of 0.4 was used to identify connected genes. *Cpt2* is the second top gene with the highest connectivities.

The results from IVCCA shows that *Cpt2* has the highest “degree” since it has the largest number of interactions with other genes above a correlation threshold of 0.75. This top-ranking gene from IVCAA is also the top gene from the SPRING database (**Figure 10B)**. *Cpt2* is highly expressed in the heart and encodes for the enzyme carnitine palmitoyltransferase 2, which is involved in the transportation of long-chain FAs into the mitochondria for energy production (73). In oxaliplatin treated mice, *Cpt2* is downregulated (FC = −1.71). Mice with *Cpt2* loss have developed cardiac hypertrophy and systolic dysfunction (74). In patients with severe *Cpt2* genetic defects, cardiomyopathy is commonly seen, and a diet enriched with medium-chain FAs is used as therapy to compensate for the reduced calorie intake from avoiding long-chain FAs (75). A similar strategy could be used for treating patients with oxaliplatin induced CTX. However, this idea has to be approached with caution. *Cpt2* shows a high negative correlation with *Trp53* (*q* = −0.92) that encodes tumor protein p53 and other genes in the cell cycle and colorectal cancer pathways (**Figure S12**). Trp53 is a tumor suppressor gene that plays a crucial role in regulating cell cycle, DNA repair, and apoptosis. Boosting the *Cpt2* expression might significantly alter p53 function that could potentially reduce the sensitivity of cancer cells to the effects of chemotherapeutics and negatively influence the effectiveness of oxaliplatin.

In contrast, genes with fewer connections (lower degree node) can also present an interest as potential therapeutic targets. These genes can be considered as downstream in the pathway. Targeting these genes might lead to fewer interactions with the other pathways and have fewer side effects. Their lower connectivity suggests they may have a more specific role and less influence on multiple pathways or processes, reducing the likelihood of disrupting other vital functions in the cell. One of these genes is *Nmrk2* that encodes enzyme nicotinamide riboside kinase 2 (NRK2) that plays a critical role in the biosynthesis of nicotinamide adenine dinucleotide (NAD^+^), which is essential for various metabolic processes in the heart, maintaining cellular homeostasis and energy balance. As shown in our previous report *Nmrk2* shows a remarkable overexpression with the FC= 16.6 and its activation can be considered as a protective mechanism to increase the lowered level of NAD^+^. *Nmrk2* has a relatively low and positive correlation with *Trp53* (q = 0.51) and relatively low correlation with the cell cycle and colorectal cancer pathways (**Figure S13**). For that matter, the entire nicotinate and nicotinamide metabolism pathway (KEGG, mmu00760) seem to be relatively safe to target given its low correlation with both the cell cycle and colorectal cancer pathway (see **Table 2**). Future experiments will be designed to test this hypothesis in cancer models in animal model systems.

### Predicting the function of unknown genes and their potential in CTX treatment

Our dataset revealed a significant portion (nearly 9%) of DEGs with unknown functions, commonly identified by prefixes such as ‘Gm’ or suffixes like ‘Rik.’ These genes often lack defined roles as yet and standard names, usually discovered via genomic sequencing and bioinformatic analysis. To predict potential functions for these genes, our approach involves two main strategies: using Gene_to_Pathways algorithm that analyzes the genes’ relationships with specific biological pathways and Gene Proximity Analysis. The latter identifies genes located near the unknown genes by examining clustering plots. **Figure 11A** illustrates this method with 10 genes closely located to an unknown gene on a t-SNE plot. This approach not only aids in understanding the unknown genes functions but also predicts their potential as suitable targets for further research.

**Figure 11.**
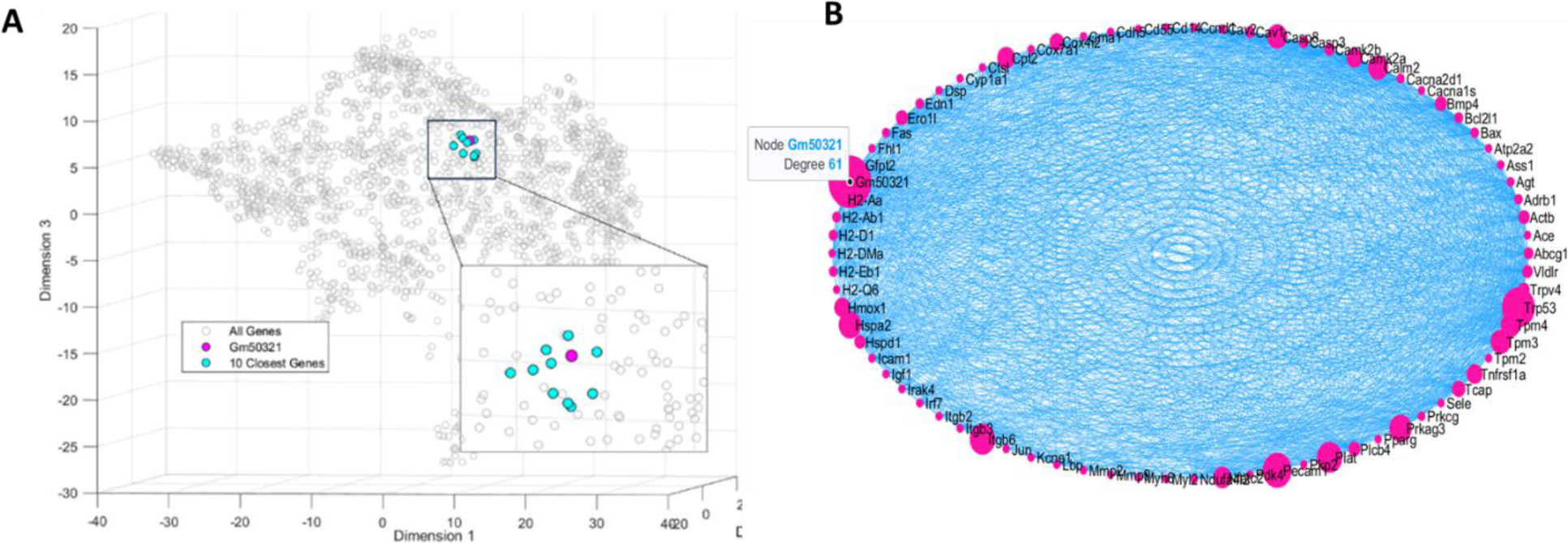
Exploring unknown gene function with close proximity analysis and network using *Gm50321* as an example. **A**: Genes identified through proximity mapping in the t-SNE plot highlights a cluster of 10 genes closely associated with *Gm50321: Cd83, Cnn2, Gm13493, Intu, Ldhd, Kdsr, Pls3, Slc25a42, Trp53, Tspan2, Zfp612* (notice the close proximity of *Trp53* to *Gm50321*). **B:** Network analysis across genes from the heart related pathways (KEGG, total number of genes 78 found among the DEGs). *Gm50321* has the highest degree among all other genes. Correlation threshold = 0.75.

For example, *Gm50321*, prominently expressed in heart tissue, showed a marked decrease in expression (FC=-1.92) in the cardiac tissues of oxaliplatin-treated animals. Analyses utilizing Gene_to_Pathways and Gene Proximity techniques indicated an association of *Gm50321* with a range of metabolic activities, muscle contraction, and mitochondrial functionalities. Notably, this gene exhibited a high correlation with cardiovascular pathways, achieving the highest average correlation (above 0.8) among all *Gmxxx* and *xxxRik* genes identified within our DEGs dataset. Network analysis incorporating cardiac function-related genes from the KEGG database and pathways identified in the literature (66–69), indicated that *Gm50321* has the highest connectivity as demonstrated in **Figure 11B**. This suggests that *Gm50321* may be a central gene regulating heart activity in oxaliplatin-treated mice.

While targeting *Gm50321* is anticipated to achieve cardiac benefits in CTX, its implications for oxaliplatin treatment and cancer management is questionable. This is primarily due to the strong negative correlation of *Gm50321* with *Trp53* (correlation coefficient, q = −0.96). Additionally, this gene exhibits substantial correlations with the cell cycle (correlation = 0.784) and colorectal cancer pathways (correlation = 0.826), suggesting a potential interference with cancer treatment. On the contrary, another gene of unknown function, *Gm31520*, demonstrates a more promising profile **Figure S12**. This gene maintains significant correlations with cardiac-related pathways, including cardiac hypertrophy (correlation = 0.780) and cardiac myopathy (correlation = 0.760 while it shows relatively lower correlations with *Trp53* (*q* = 0.596), the cell cycle (correlation = 0.484), and colorectal cancer pathway (correlation = 0.350). Close proximity genes pointed to the tight relationship with the genes involved in the assembly of actin filaments (via *Pdlim3*) and a complement pathway involved in inflammation (via *C2*) (**Figure S12**). These characteristics render *Gm31520* a more promising target for mitigating CTX-induced cardiac complications without adversely affecting cancer treatment efficacy.

### Limitations of the global correlation approach

Performing a global correlation analysis considering the intervariability between the variables promises to provide insights on the biological processes and identify new pathways and targets as we have demonstrated on oxaliplatin induced CTX. This approach simplifies the analysis by condensing multiple gene-gene relationships within a dataset of all DEGs into a few metrics. This approach makes it easier to identify potential targets that avoids silencing the pathways that chemotherapy uses to destroy cancer cells or activating the pathways that cancer cells use to proliferate.

With many opportunities this approach offers, this approach has a number of weaknesses. The correlation does not imply function and the method does not take into account the physiology role of a gene in its pathway. A high correlation of the pathway to other pathways does not necessarily mean the pathway is more important. It might indicate that the genes in that pathway are more synchronized in responding to the stimuli or share a similar expression pattern under the studied conditions. The limitation of our approach lies in the heterogeneity within pathways since biological pathways often consist of genes with diverse functions and roles. If there are outlier genes within a pathway that have strong correlations with other genes outside the pathway, averaging can be influenced by these outliers and may not accurately reflect the pathway’s intrinsic coordination. Another limitation in our method lies in the treating positive and negative correlations equally, potentially missing important information. In addition, the genes with very high and very low expressions are taking equal, that might lead to the errors. An improvement can address clustering. Enhancing the clustering process in t-SNE can significantly refine data visualization and interpretation. Currently, the clustering largely depends on visual analysis without prior information about the number of clusters, which is largely subjective. A more robust, intuitive, and automated process for identifying and validating clusters in high-dimensional data would significantly improve the data analysis. Finally, the proposed method can be used for relatively large number of DEGs such as in this example where we have 1744 DEGs. Smaller number, for example less than a few hundred, might not provide sufficient data to build a comprehensive visualization of connectivity between the genes.

## SUMMARY

The Inter Variability Cross-Correlation Analysis (IVCCA) platform offers a powerful method for analyzing RNA-seq and other high-throughput data, unveiling layers of information not previously accessible. This approach was tested on RNA-seq data from the heart tissues of mice treated with oxaliplatin. The methodology ranked genes by their correlation to other differentially expressed genes, identifying important genes and pathways. Using this method, we have identified genes and pathways that are central to oxaliplatin induced CTX and pathways that can be targeted to potentially to minimized adverse effects. Notably, this includes metabolic energy pathways, which are a safe target for mitigating cardiotoxic side effects due to their minimal interaction with oxaliplatin’s cancer-targeting pathways. Furthermore, the IVCCA approach has uncovered potential inter-organ associations, such as between the kidney and the heart. Future developments will include validation of our finding through synthetic biology and genetic engineering and targeting specific genes for therapeutic intervention. In addition to developing new drugs, this approach might lead to repurposing opportunities by linking existing drug action mechanisms with identified gene correlation networks.

## Supporting information

All_Genes

All_DEGs

Random_Selected_Genes

## ACKNOWLEDGMENTS

MB acknowledges support from the NIH (R01 CA208623 and R21CA269099).

## SUPPLEMENTAL INFORMATION

1. Theory of IVCCA analysis

2. Lists of all genes, DEGs, randomly selected genes and their expressions

3. Lists of pathway-specific genes

4. MATLAB code for IVCCA: https://github.com/MikhailBerezin/IVCCA/

