## SUPPLEMENTAL INFORMATION for "Identification Drug Targets for Oxaliplatin-Induced Cardiotoxicity without Affecting Cancer Treatment through Inter Variability Cross-Correlation Analysis (IVCCA)"

**Institutional Affiliations**: ^a^Mallinckrodt Institute of Radiology, Washington University School of Medicine St. Louis, MO 63110, USA; ^b^Institute of Materials Science & Engineering Washington University, St. Louis, MO 63130, USA; ^c^Cardio-Oncology Center of Excellence, Washington University School of Medicine, St. Louis, MO 63110; Department of Genetics, Washington University School of Medicine, St. Louis, MO 63110

### THEORY

##### Pairwise Pearson correlation between two genes

The Pearson correlation coefficient of two variables (genes) is a measure of their linear dependence. For each gene of interest with n scalar members (in our case the *n* = number of mice), the Pearson correlation coefficient is defined by Eq.1:

| $q\left( A,B \right)=\frac{1}{n}\sum_{i=1}^{n} \left( \frac{A_{i}-\mu_{A}}{\sigma_{A}} \right)\left( \frac{B_{i}-\mu_{B}}{\sigma_{B}} \right)$ | 1 |
| --- | --- |

where μA and σA are the mean and standard deviation of the randomly selected gene A, respectively, and μB and σB are the mean and standard deviation of the randomly selected gene B.

Pearson correlation coefficient values lie in the range between -1 and +1, where values close to +1 indicate a strong positive linear relationship between the two genes being compared, while values close to -1 suggest a strong negative linear relationship. A Pearson correlation coefficient of 0, implies no linear relationship or correlation between the genes.

##### Global correlation analysis

The global correlation analysis involves calculating the average of the sum of absolute Pearson correlation coefficients within a dataset of genes. For a dataset of DEGs with *n* genes, denoted as *G_1_, G_2_, …, G_n_* each gene is measured across *m* samples where *m* is the number of mice:

| $G_{i}=\left\{ G_{i1} ,G_{i2} ,\ldots,G_{im} \right\}$ | 2 |
| --- | --- |

For each gene *G_i_*, the sum of absolute Pearson coefficients $q$ with all other genes *G_j_* ($i\neq j$) can be calculated from Eq. 3:

| $G_{i}= \sum_{j\neq i} \vert q\left( G_{i},G_{j} \right)\vert$ | 3 |
| --- | --- |

where *G_i_* – is the sum of absolute correlation coefficients of the gene *i* calculated as to the sum of absolute correlation coefficients $q_{i}$ for all pairwise genes in the dataset relative to this gene.

The average of these sums of absolute correlations across all *n-1* genes in the dataset while ignoring self-correlations can be expressed by Eq. 4:

| $Q_{i}=\frac{G_{i}}{n-1}= \frac{1}{n-1}\sum_{j\neq i} \vert q\left( G_{i},G_{j} \right)\vert$ | 4 |
| --- | --- |

The result of this equation provides a numerical value that summarizes the correlation structure for each gene within the gene dataset. The global correlation value *Q_i_* indicates how closely related the genes are on average across the entire dataset.

*Q* values lie within a range of 0 to 1. Higher values indicate a stronger overall correlation to the genes in the dataset with the value of 1 indicating a perfect positive correlation among all genes.

##### Pathway Correlation Indices (PCI_A and PCI_B)

The correlation value for the entire pathway is defined by the Pathway Correlation Indices (PCI_A and PCI_B). The difference between *PCI_A* and *PCI_B* is that *PCI_A* is calculated by averaging individual absolute correlation values $\tilde{Q}_{i}$with all genes within the pathway, while *PCI_B* is calculated by averaging the extracted absolute correlation values from the entire dataset as shown in **Figure S2** according to Eq. 5 :

| $PCI\_A=\frac{1}{L}\sum_{i=1}^{L} \tilde{Q}_{i}; PCI\_B=\frac{1}{L}\sum_{i=1}^{L} Q_{i}$ | 5 |
| --- | --- |

where *L* is the number of DEGs in the specific pathway, $\tilde{Q}_{i}$ – the correlation values between genes within the pathway, $Q_{i}$is the correlation values between genes from the pathway to all genes in the dataset. In the extreme case where the number of genes in the pathway equal to the number of genes in the entire dataset *PCI_A = PCI_B*

A PCI_A value indicates how tight the genes within the pathway correlate to each other, while a PCI_B value indicates how strong the impact of the genes from the pathway to the entire dataset. The latter value can be used to rank the pathways as well as compare two or more pathways.

**Note 1**: large number of DEGs might be computationally demanding, since the total number of unique pairwise *k*=2 combinations of *n* genes can be estimated according to the **Eq. 6**:

| $C\left( n,k \right)=\frac{n!}{k!\left( n-k \right)!}= \frac{n!}{2\left( n-2 \right)!}$ | 6 |
| --- | --- |

**Note 2**: This approach treats positive and negative correlations equally, which may not be appropriate for some specific pathways.

*Kullback-Leibler Divergence*: We used Kullback-Leibler Divergence to calculate the difference between two joint probability distributions derived from the original correlation matrix (*Pm*) and from the t-SNE data (*Qm*). The method involves several steps: i) joint probability distribution creation, ii) normalization of the joint probability distribution and iii) calculating the KL divergence. KL values closer to the zero are considered to be ideal. The details of the calculation are given in the Supplemental information.

Firstly, for the original correlation matrix, the correlation scores are transformed into distances using the **Eq. 7:**

| $distances=1-abs(data)$ | 7 |
| --- | --- |

These distances are then transformed into similarities using a Gaussian kernel. The spread of the Gaussian kernel is controlled by the parameter sigma, which we set equal to the perplexity value in our calculations. The similarities are calculated as:

| $similarities= \exp\left[ \frac{-distances}{2\times{sigma}^{2}} \right]$ | 8 |
| --- | --- |

These similarities are then normalized row-wise to convert them into conditional probabilities. The conditional probabilities are symmetrized to form the joint probability matrix, *P_m_*, for the original high-dimensional data and normalized. Normalization is performed by dividing each element by the sum of all elements in the distribution. This scaling transforms the raw scores into a range that represents probabilities where each value is between 0 and 1, and the sum of all values is 1.

| $P_{m}=\frac{P_{m}}{\sum P_{m}}$ | 9 |
| --- | --- |

Similarly, the t-SNE joint probability distribution (*Q_m_*) is calculated based on the low-dimensional representation obtained through t-SNE. It first calculates pairwise Euclidean distances for the t-SNE representation of the data. These distances are then converted into similarities using the same sigma value and transformed into conditional probabilities. The conditional probabilities are symmetrized to obtain the joint probability distribution, *Q_m_*, representing the t-SNE Result Distribution and normalized in a:

| $Q_{m}=\frac{Q_{m}}{\sum Q_{m}}$ | 10 |
| --- | --- |

Finally, the KL Divergence is calculated by summing the product of the elements of Pm and the logarithm of the ratio of *P_m_* to *Q_m_*, over all non-zero elements of *P_m_*.

| $KL=\sum P_{m}\times log \frac{P_{m}}{Q_{m}}$ | 11 |
| --- | --- |

Overall, this divergence measures how much the probability distribution over the original data (*P_m_*) diverges from the probability distribution over the low-dimensional representation (*Q_m_*), providing a numerical value of how well t-SNE algorithm is performed in terms of being close to the original data. KL values closer to the zero are considered to be ideal.

##### Comparison of two pathways with cosine similarity.

The algorithm first finds overlapping genes between two pathways among the genes in the dataset. The algorithm then calculates cosine similarity using **Eq. 12-13**.

| $Cosine similarity=\frac{Dot\_product}{\sqrt{Norm\_set1}\times\sqrt{Norm\_set2}}$ | 12 |
| --- | --- |

Where *Norm_set1* and *Norm_set2* are the counts of unique genes in each pathway plus the number of overlapping genes.

| $Dot\_product=\sum_{i=1}^{N} {q_{i}}^{2}+ \sum_{j=1}^{M} 1$ | 13 |
| --- | --- |

*where Dot_product* is the sum of squared correlation values for each non-overlapping gene pair extracted from the entire correlation matrix with absolute values, plus 1 for each overlapping gene. *N* is the number of non-overlapping gene pairs. *q_i_* represents the value associated with the *i*-th non-overlapping gene pair; *M* is the number of overlapping genes.

##### Multiple pathways analysis

The function calculates the following metrics:

*Pathway Activated Index: PAI* (defined as Eq. 14)*,*

*Pathway Correlation Indices (PCI_A and PCI_B),*

*Correlation-Expression Composite Index (CECI* (Eq. 15)*,*

*Z-score* (Eq. 16)*,*

*Z-score-critical* (Eq. 17)*.*

*PAI* quantifies the extent to which a particular pathway is activated and is defined as the ratio of differentially expressed genes found in the set to the total number of genes in the pathway.

| $PAI= \frac{Differentially expressed genes from the pathway}{Total number of genes in the pathway}$ | 14 |
| --- | --- |

| $CECI=PAI \times PCI_{B}\times100$ | 15 |
| --- | --- |

*CECI* is a composite measure that reflects both the correlation among genes within the entire dataset (as indicated by *PCI_B*) and the proportion of DEGs in that pathway.

The Z-score provides a statistical measure of how different a given pathway is from a normative dataset composed of pseudo-pathways. Pseudo-pathways were generated from random sets of 50 to 200 genes which is a typical size of the pathway such as KEGG and GO from all genes in the set (13775 genes in the case used in this study). This process was repeated 100 times resulting in 100 different pseudo-pathways. *CECI* values for each pseudo-pathway (=*Y*) were calculated using Multiple pathways analysis tool (see above). The average *Average(Y)* and standard deviation *STD(Y)* of these values were found to be equal to 7.91 and 2.061 correspondingly. The *Z-score* was then calculated by summing the absolute correlations for each gene with other genes in the set, then averaging these sums across all genes in the set:

| $Z\_score=\frac{X-Average(Y)}{STD (Y)}$ | 16 |
| --- | --- |

where for *X* equal to the CECI value of a pathway, *Average (Y)* = 7.91 and *STD (Y)* = 2.0605.

To determine statistical significance, we calculated the *Z_score_critical* value (Eq. 17). We choose a significance level α=0.05. For a two-tailed test, the *Z_score_critical* is approximately 1.96. Pathways with the *Z_score* > *Z_score_critical* were considered statistically significant and graphically presented as bar graphs.

| $Z_{score_{critical}}=Ф^{-1}\left( 1-\frac{\alpha}{2} \right)=1.96$ | 17 |
| --- | --- |

Where $Ф^{-1}$ is the inverse of the standard normal cumulative distribution function, α is the significance level (e.g., 0.05 for a 5% significance level).

##### Network analysis

The function retrieves gene correlation data and gene names and prompts the user to enter a correlation threshold. This threshold is used to filter the correlation data, keeping only the gene pairs that exhibit a correlation higher than a threshold. The function constructs a 3D network graph, where each node represents a gene, and edges connect genes with correlations exceeding the threshold. It calculates the '*degree*' of each gene, which is the number of connections to other genes, indicating its connectedness within the network. The results are displayed in a user interface as a sortable table, listing each gene alongside its degree. For better visualizing we used a node size enhancement formula (Eq. 17).

| $Node_{degree}=5+ \left[ \frac{1.3*(Node_{degree})}{max(Node_{degree})} \right]^{n}$ | 18 |
| --- | --- |

where *Node_degree_* is the degree of the node, and *max(Node_degree_)* is the maximum degree among all nodes in the network, *n* – is the enhancer that can be used between 6 and 12

To visually map the strength of correlations the thickness of edges in the graph were scaled to ensure that all lines have a minimum visibility and that more substantial correlations are increasingly emphasized. Eq. 18 scales the size of each node relative to the maximum degree, ensuring that nodes with more connections are visually larger. The constants 5 and 1.3 and the power of 12 are used to amplify the differences in size, making it easier to distinguish between nodes with varying degrees of connectivity.

| ${edge}_{thickness}=minLineWidth+ \left[ \frac{EdgesWeight}{maxWeight}\times0.32(maxLineWidth-minLineWidth \right]^{m}$ | 19 |
| --- | --- |

Where *maxWeight* is the maximum weight among all the edges in the graph. The weight represents the strength or magnitude of the correlation between nodes in the graph. The maximum value is used as a reference point to scale all other weights. *minLineWidth* is the desired min line thickness on the graph, set to 0.5, *maxLineWidth* is the desired max line thickness, set to 4. *m* – is the enhancer that can be used between 6 and 12.

### FIGURES

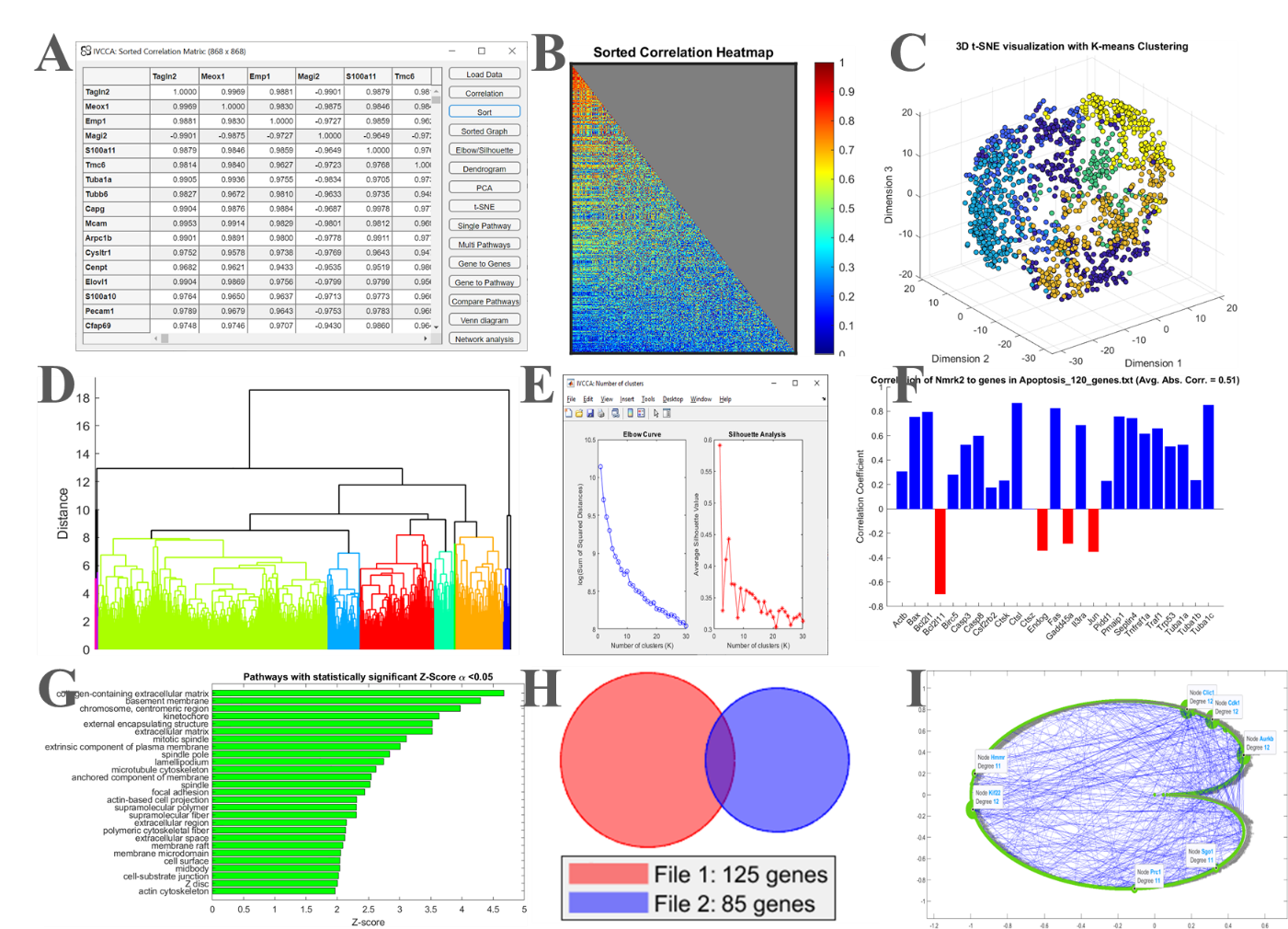

**Figure S1** **The** **IVCAA GUI of the Correlation Analysis tool**. **A.** Loading the data, where each column represents a different gene, while each raw represent the sample. **B.** Visualization of the sorted correlation matrix **C**. Clustering toolboxes (shown t-SNE); **D**. Color coded dendrogram with clustering **E**. Elbow/Silhouette estimating the number of list of clusters. **F.** Correlation of a gene to a group of genes dendrogram showing clusters. **G**. Top statistically significant pathways. **H.** Venn diagram of two pathways **I.** Network analysis. Available for download in MATLAB or an executable file per request.

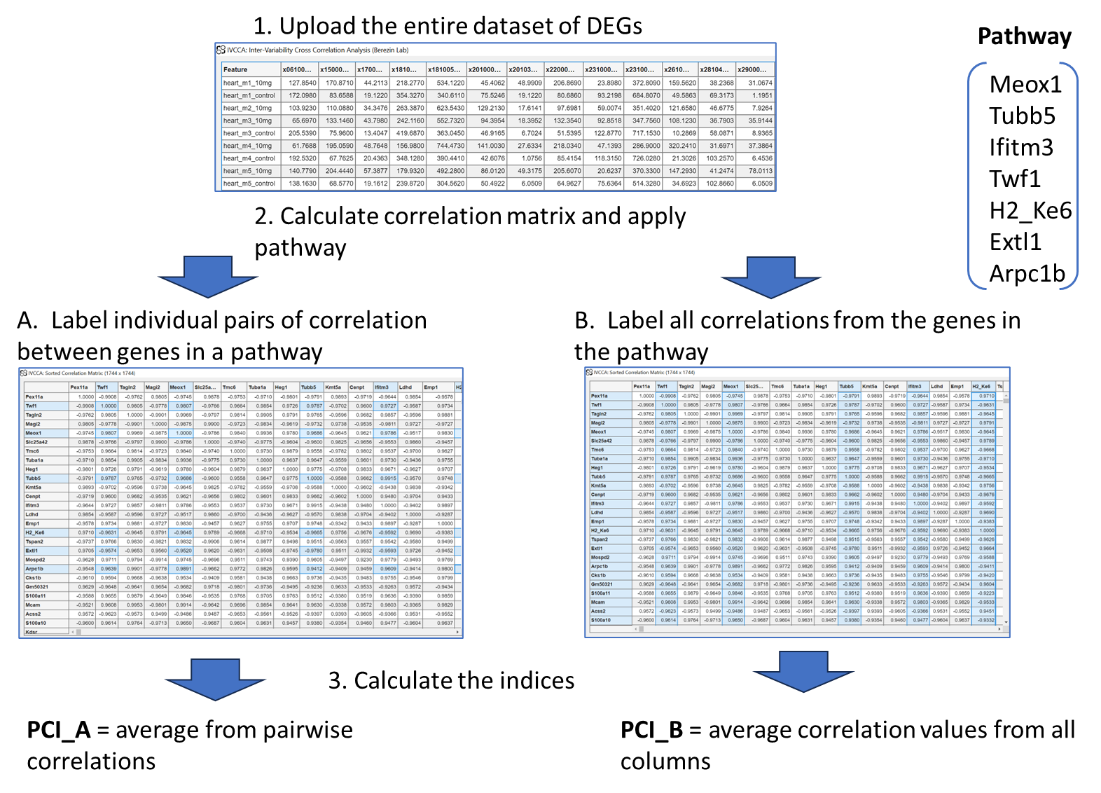

**Figure S2** Illustration the difference between *PCI_A* and *PCI_B* is that *PCI_A* is calculated by averaging individual absolute correlation values $\tilde{Q}_{i}$with all genes within the pathway, while *PCI_B* is calculated by averaging the extracted absolute correlation values from the entire dataset.

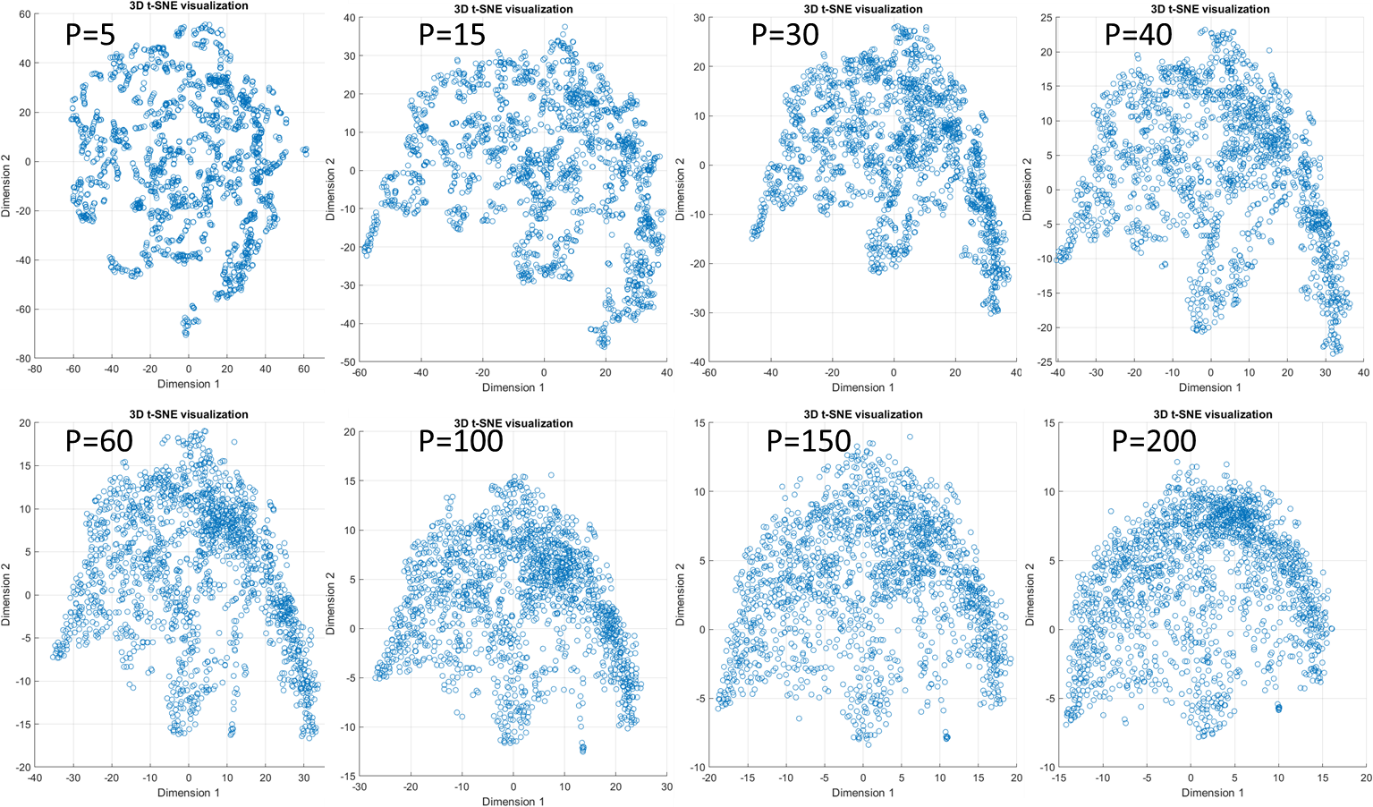

**Figure S3** t-SNE plots of the correlation matrix at different values of perplexity (P). Visual evaluation of a 2D view suggests P=60 as the most optimal perplexity

| Perplexity | KL divieregnece |
| --- | --- |
| 5 | 0.002637 |
| 15 | 0.004229 |
| 30 | 0.0035189 |
| 40 | 0.0011163 |
| 60 | 0.00018097 |
| 100 | 8.3781e-06 |
| 150 | 5.2615e-07 |
| 200 | 7.326e-08 |

**Figure S4** Effect of perplexity on KL divergence.

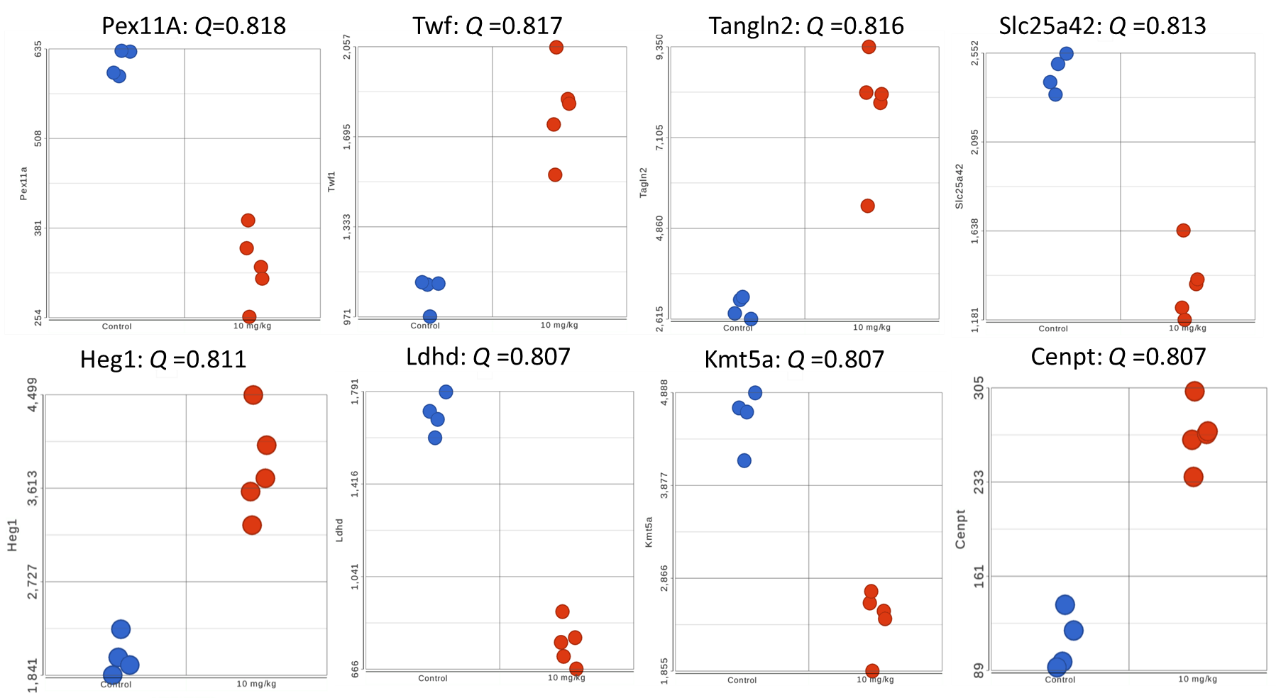

**Figure S5** Change in the expression of the selected top-ranking genes from the cross-correlation analysis of 1744 DEGs (FDR<0.05, FC>1.5). Q is defined as an average Pearson coefficient across the entire gene dataset

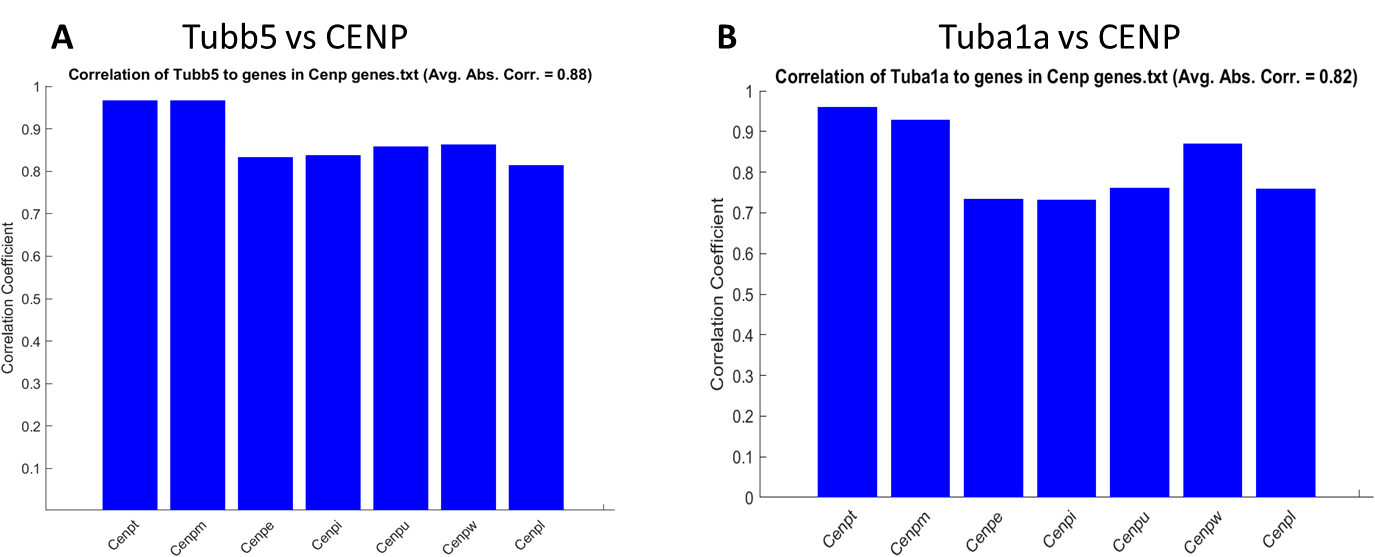

**Figure S6** Correlation between tubulin encoded genes and centromere protein expressing genes

Statistically significant pathways from the KEGG enrichment analysis

|  | Enrichment score |
| --- | --- |
| Complement and coagulation cascades | 22.7467 |
| Neuroactive ligand-receptor interaction | 17.8535 |
| Staphylococcus aureus infection | 15.952 |
| p53 signaling pathway | 9.96626 |
| Pathways in cancer | 8.73952 |
| Systemic lupus erythematosus | 8.45305 |
| Calcium signaling pathway | 8.26251 |
| Cytokine-cytokine receptor interaction | 7.64893 |
| Vascular smooth muscle contraction | 7.38103 |
| Small cell lung cancer | 7.27303 |
| Cell adhesion molecules | 7.27303 |
| Amoebiasis | 7.24166 |
| Steroid hormone biosynthesis | 7.04123 |
| Protein digestion and absorption | 6.89743 |
| Hematopoietic cell lineage | 6.89743 |
| Phagosome | 6.79379 |
| Pertussis | 6.66959 |
| Renin-angiotensin system | 6.5788 |
| Viral myocarditis | 6.48422 |
| Viral protein interaction with cytokine and cytokine receptor | 6.07011 |
| Transcriptional misregulation in cancer | 5.83005 |

**Figure S7** KEGG enrichment analysis performed using Partek Flow software. The pathways were filtered for FDR<0.05.

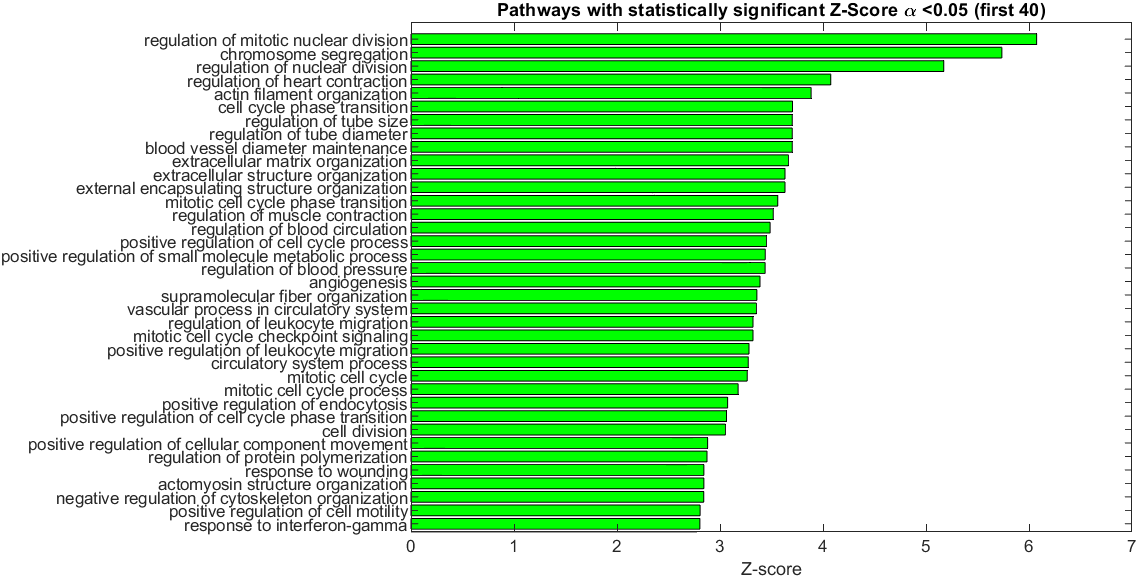

**Figure S8** **GO biological functions**: top 40 ranked biological function out of a total 128 significant pathways. The top biological functions are related to the cell cycle, regulation of blood circulation, and regulation of the metabolic processes.

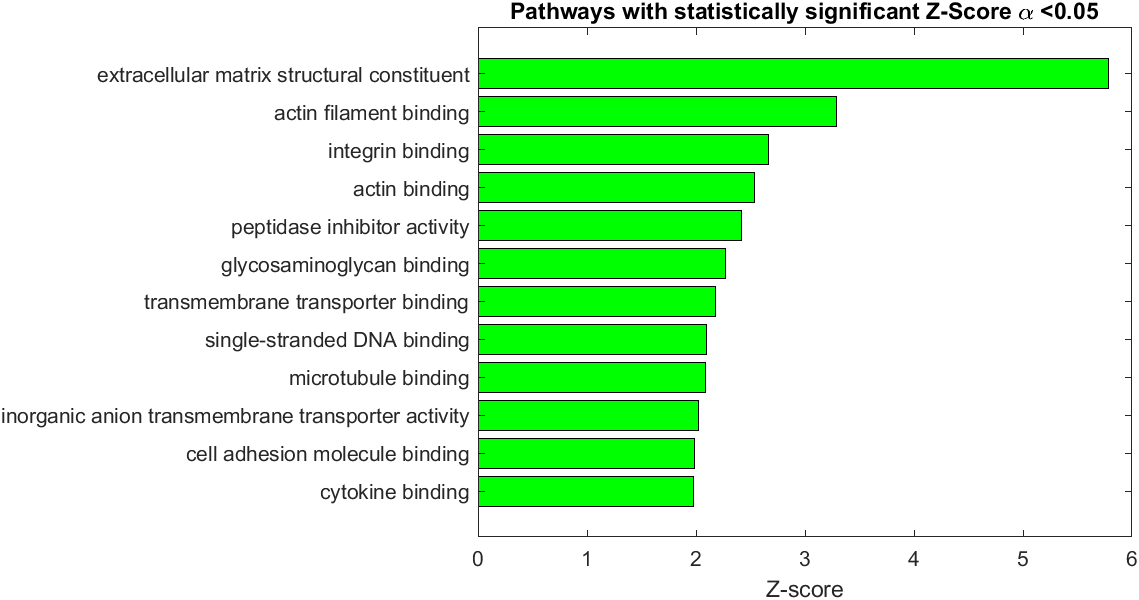

**Figure S9** **GO molecular functions**: top ranking molecular function: 12 significant pathways. The top molecular functions are related to extracellular matrix processes, binding to actin, and transmembrane transport.

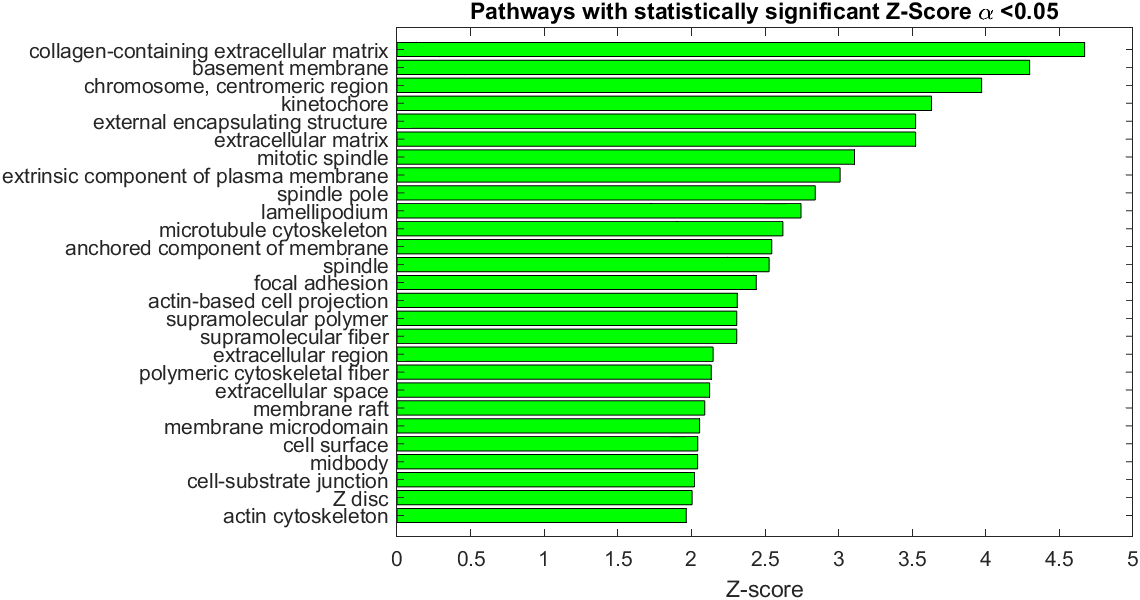

**Figure S10** **GO cell components**: 27 significant pathways. The top cell components are related to extracellular matrix and chromosomes.

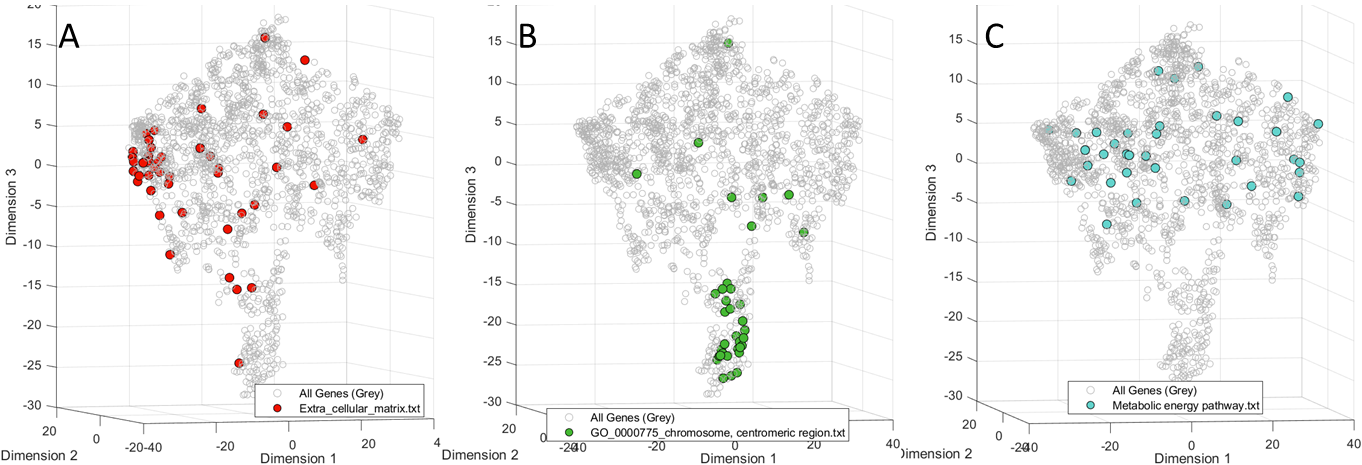

**Figure S11** Representative projections of pathways on the t-SNE 3D scatter plot. **A**: Extracellular matrix (1), **B**: GO_0000775 Chromosome centromeric region pathway, **C**: metabolic energy pathway (2)

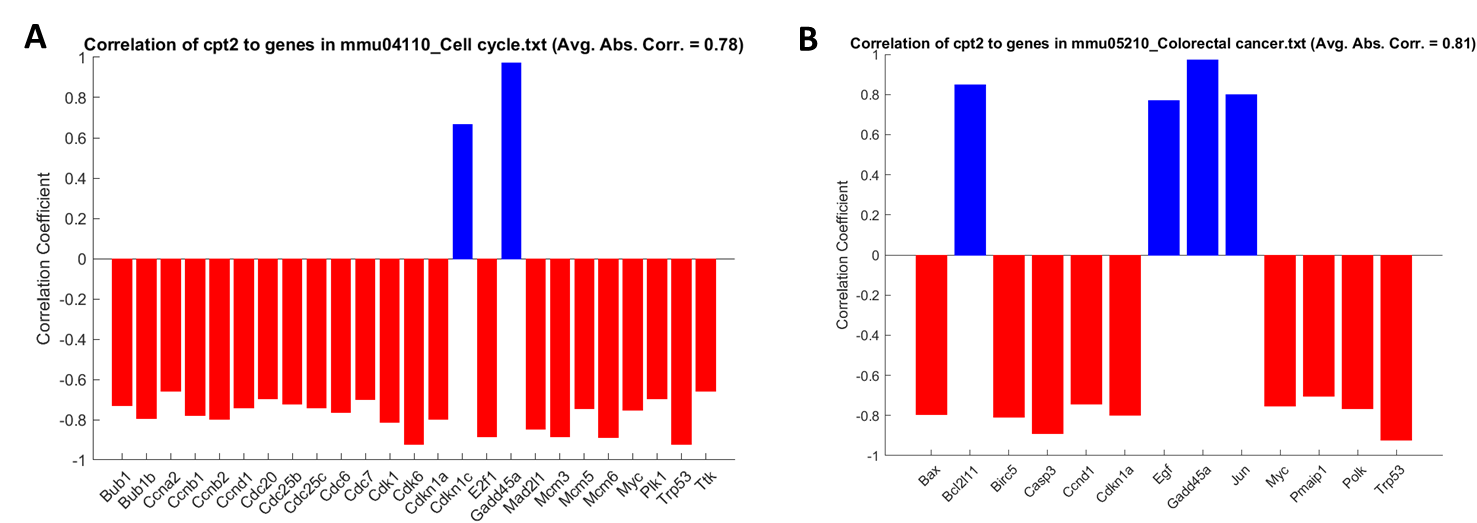

**Figure S13** Correlation of *Cpt2* to individual genes in **A**: cell cycle pathway and **B**: colorectal cancer pathway. The correlations are high for for most of the genes with the average absolute correlation 0.78 for the cell cycle and 0.81 for the Colorectal Cancer

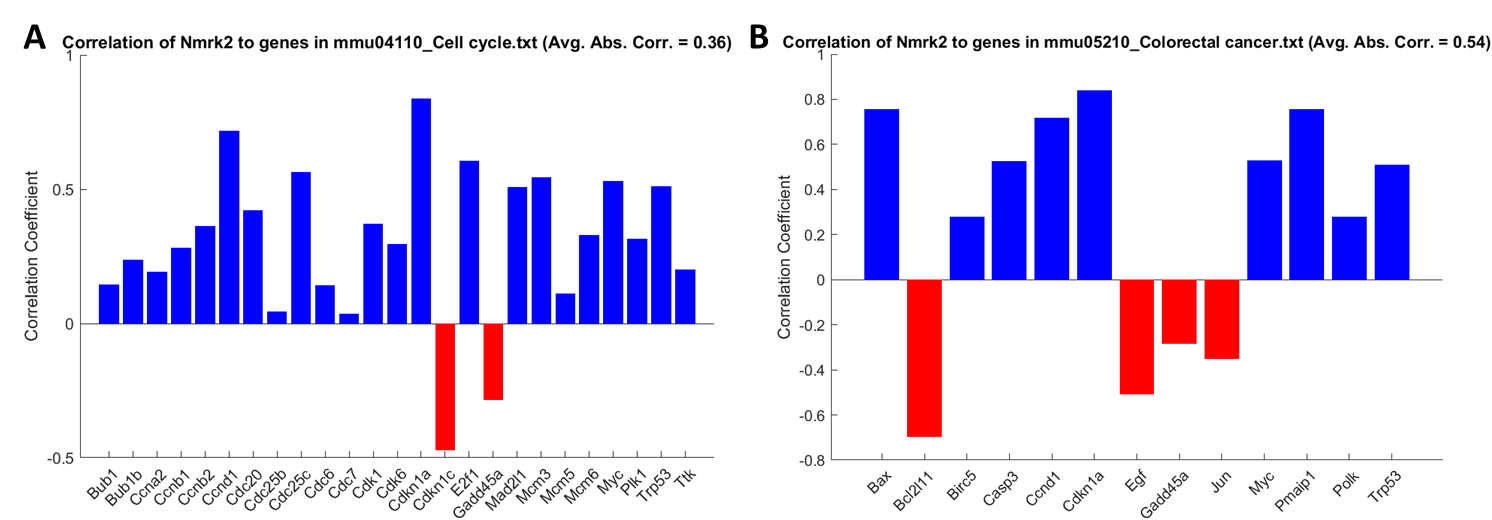

**Figure 14** Correlation of *Nmrk2* to individual genes in **A**: cell cycle pathway and **B**: colorectal cancer pathway. The correlations are in general low.

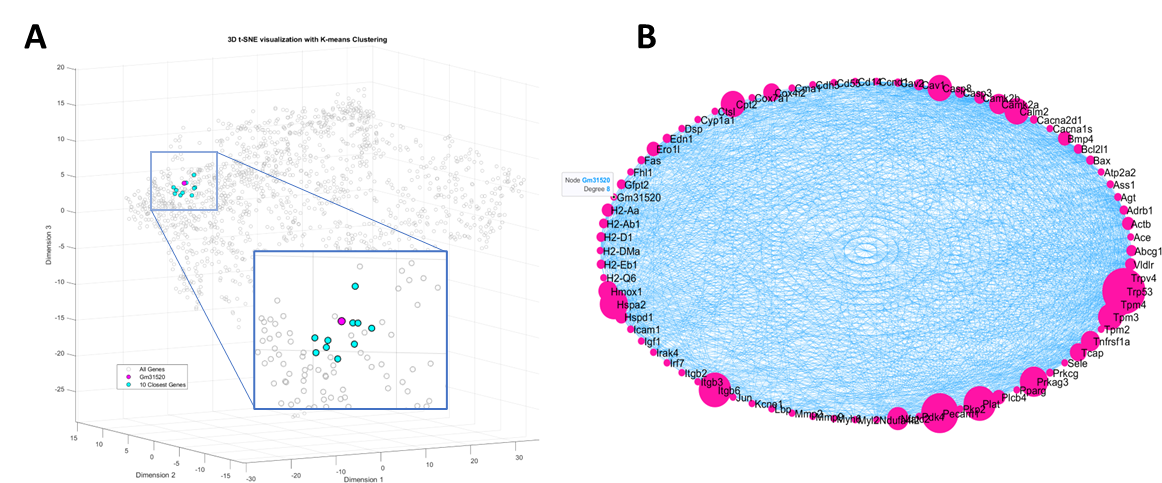

**Figure S12** C**lose proximity analysis of Gm31520.** **A**: Genes identified through proximity mapping in the t-SNE plot highlights a cluster of 10 genes closely associated with Gm31520: Ctla2a, Chn1, Cy2s1, Msln, Tat, Pdlim3, Slc12a8, C2, Golm1, Fam114a1. **B:** Network analysis across genes from the heart related pathways (KEGG, total number of genes 78 found among the DEGs). Gm50321 has the highest degree among all other genes. Correlation threshold = 0.75

**Table S1 Genes closest to** **Gm50321 from the close proximity analysis and their relevant function**

| **Gene** | **Fold change** | **Heart related function** | **Ref** |
| --- | --- | --- | --- |
| Cd83 | -3.65 | Immune function, dendritic and B cells | (3) |
| Cnn2* | 1.72 | Muscle contraction regulation | (4) |
| Ldhd* | -2.20 | Lactate metabolism | (5) |
| Intu | -1.80 | Cell signaling and organization (largely unknown) | (6) |
| Kdsr | 1.67 | Sphingolipids synthesis, detoxification | (7) |
| Pls3 | 1.91* | Actin filament organization | (8) |
| Slc25a42* | -1.79 | Mitochondrial function and energy metabolism | (9) |
| Trp53* | 1.56 | Cellular stress, apoptosis, ROS | (10) |
| Tspan2 | 2.64 | Cell signaling, blood vessels | (11) |
| Zfp612 | -1.74 | Predicted transcription factor (largely unknown) | - |

*) highly expressed

**Table S1 Genes closest to Gm31520 from the close proximity analysis and their relevant function**

| **Gene** | **Fold change** | **Heart related function** | **Ref** |
| --- | --- | --- | --- |
| Ctla2a* | 2.77 | Expressed in activated T-cells, secretory factor | (12) |
| Chn1 | 1.58 | Neuronal signal-transduction mechanisms | (13) |
| Cyp2s1 | 4.79 | Oxidative metabolism, metabolism of toxic compounds | (14) |
| Msln** | 11.09 | Mesothelin repair, cellular adhesion | (15) |
| Tat | 7.92 | Transferase that interconverts tyrosine and glutamate, norepinephrine synthesis | (16) |
| Pdlim3* | 1.77 | Actin filament in muscle cells | (17) |
| Slc12a8 | 2.84 | NAD+ transport | (18) |
| C2 | 3.47 | Part of the complement system, inflammation | (19) |
| Golm1 | 1.59 | Response protein to viral infection | (20) |
| Fam114a1 | 1.81 | Heart remodeling via angiotensin II | (21) |

*) highly expressed

**) become highly expressed after oxaliplatin
